# Co-targeting MERTK and EGFR with a Bispecific Antibody Overcomes Drug Resistance Across Mutations in Exons 19, 20, and 21

**DOI:** 10.1101/2025.10.16.681972

**Authors:** Suvendu Giri, Boobash-Raj Selvadurai, Arturo Simoni-Nieves, Nitin Gupta, Moshit Lindzen, Rishita Chatterjee, Alessandro Genna, Marieke Van Daele, Deepthi Ramesh-Kumar, Mirie Zerbib, Roni Oren, Isabelle Sophie Meijer, Elisabeth Franziska Wippich, Yahel Avraham, Rony Dahan, Laura E. Kilpatrick, Simon Platt, Stephen J. Hill, Lipika R. Pal, Eytan Ruppin, Donatella Romaniello, Mattia Lauriola, Yosef Yarden

**Author notes:** To whom correspondence should be addressed: Yosef Yarden, Department of Immunology and Regenerative Biology, Weizmann Institute of Science, Rehovot 76100, Israel. Department of Life Science - Molecular Biology, University of Vienna, Wien, Austria (Isabelle Sophie Meijer). **Declaration of interests:** All authors declare that they have no conflict of interest relevant to the herein reported study.

## Abstract

Resistance of lung cancer to EGFR-specific tyrosine kinase inhibitors (TKIs), such as osimertinib, often arises from secondary mutations or the activation of bypass signaling pathways. We noted elevated levels of GAS6 and its receptor, MERTK, in patients acquiring resistance to osimertinib, hence hypothesized that the GAS6–MERTK axis can serve as a therapeutic target. GAS6 promoted cell survival and an anti-GAS6 antibody we generated delayed tumor relapses in a xenograft model. Likewise, MERTK ablation sensitized lung cancer cells to osimertinib due to a newly described EGFR-to-MERTK crosstalk. Hence, we developed Bis3, a bispecific antibody targeting both MERTK and EGFR and promoting their degradation. Bis3 plus TKIs cooperatively inhibited the growth of TKI-resistant lung cancer spheroids and markedly delayed relapses of patient-derived xenografts harboring the clinically challenging exon-20 mutations. These findings establish the GAS6–MERTK axis as a driver of drug resistance and provide a rationale for clinical development of Bis3.

## Introduction

The American Cancer Moonshot initiative, which aimed at improving cancer therapy, recommended that overcoming drug resistance be considered one of the top priorities in oncology ^1^. Indeed, the majority of patients with advanced cancer die either because their cancers are inherently resistant to anti-cancer drugs, or their cancers initially respond to specific drugs but they later develop tolerance ^2^. Resistance universally limits application of not only chemotherapeutic agents but also kinase inhibitors ^3^, anti-receptor antibodies ^4^ and immune checkpoint inhibitors ^5^. Lung cancer provides an example relevant to the importance of understanding and overcoming resistance to anti-cancer drugs. This malignancy is the leading cause of oncology-related deaths and over 75% of all cases are classified as NSCLC (non-small cell lung cancer). A significant fraction of NSCLC patients present activating mutations in the epidermal growth factor receptor (EGFR) ^6–10^. The most common mutations are exon-19 deletions and an exon-21 point mutation, L858R ^11^.

The first-generation EGFR-specific TKIs achieved superiority in comparison to chemotherapy, in terms of progression-free survival ^12–14^. However, despite initial responses, patients inevitably become resistant within 10-20 months. The most common mechanism of resistance involves emergence of the T790M secondary mutation, which increases EGFR’s affinity toward ATP ^15^. Other mechanisms include adaptive up-regulation of *MET* ^16^, *HER2* ^17^ and AXL ^18^, mutations in *RAS* ^19^ and *BRAF* ^20^, as well as phenotypic alterations ^21^. The second-generation TKIs (e.g., afatinib) irreversibly bind with the four EGFR/ERBB family members. Likewise, a third-generation TKI, osimertinib, which is able to inhibit EGFR-T790M, has been initially approved as a second-line treatment ^22^, but it was later approved as a first-line treatment ^23^. Acquired resistance limits also the application of osimertinib. The most common mechanisms of resistance to osimertinib in the first-line settings are *MET* amplification, C797X mutations, amplification of wild-type *EGFR* or *HER2,* and mutations in downstream signaling proteins ^24^.

Following treatment with TKIs, a small subpopulation of NSCLC cells harboring EGFR mutations exhibit reversible tolerance ^25,26^. However, this subpopulation of drug-tolerant persister cells (DTPs) subsequently acquires irreversible resistance. How DTP states are replaced by permanent drug-resistant states remains unclear. According to one model, TKI treated cells transiently increase their mutation rates to gain resistance ^27^. Unlike the slowly evolving pioneer mutations, which were linked to pre-existing EGFR mutations in healthy tissues ^28^, emergence of the secondary (e.g., T790M) and tertiary mutations (e.g., C797S) in patients treated with TKIs is faster and might be adaptive. Recently, two types of mutagenic mechanisms have been reported. One pathway engages apolipoprotein B messenger RNA editing catalytic polypeptide-like (APOBEC) cytidine deaminases ^29–32^. According to another report, lung cancer cells exposed to TKIs are unable to repair DNA lesions and they elevate low-fidelity (mutagenic) DNA polymerases ^33^. This second mechanism of induced mutagenesis is preceded by upregulation of GAS6 and ablation of at least one of GAS6’s receptors, AXL ^33^.

AXL, together with MERTK and TYRO3, forms the TAM family of receptor tyrosine kinases ^34^. Rather than functioning as oncogenic drivers, TAM receptors primarily promote stemness and motility ^35^. Their shared ligand, GAS6, binds phosphatidylserine exposed on apoptotic cells ^36^, enabling AXL and MERTK to act as cell death sensors. AXL is a well-established mediator of EGFR inhibitor resistance: its expression predicts poor prognosis ^37^, it supports survival of drug-tolerant persisters ^38^, and its kinase activity confers resistance ^39^. By contrast, the role of MERTK in drug resistance has been less defined. Preclinical studies showed that osimertinib resistance coincides with MERTK induction, and dual targeting of MERTK and EGFR, using the respective TKIs, reversed resistance in a tumor model ^40^. Because our re-analysis of patient data following osimertinib treatment uncovered increased expression of both MERTK and GAS6, we hypothesized that blocking either protein would suppress TKI resistance. By generating anti-GAS6 and anti-MERTK antibodies, as well as a bispecific antibody targeting both MERTK and EGFR, we demonstrate that the GAS6–MERTK axis acts as a critical mediator of drug resistance. Blocking this axis sensitizes to TKIs a wide spectrum of EGFR mutations, including not only L858R, T790M and exon-19 deletions, but also the clinically challenging exon-20 insertions. Importantly, combining MERTK and EGFR antibodies produced greater efficacy than pairing their respective TKIs. These insights, together with the antibody tools we developed, may help pave the way toward clinical translation.

## Results

### Osimertinib treatment leads to upregulation of GAS6 in patients with residual disease, but osimertinib resistance can be inhibited in animals using a new anti-GAS6 antibody

In our previous study, we demonstrated that, when applied in vitro, EGFR kinase inhibition strongly induces GAS6 expression in proliferating persister cells compared with non-cycling, drug-tolerant persisters ^33^. To assess clinical relevance, we re-analyzed single-cell RNA-seq datasets from a Phase II osimertinib trial, NCT03433469 (30 patients). Importantly, GAS6 expression was relatively low in treatment-naïve (TN) tumors, but markedly elevated in residual disease (RD), where MERTK and EGFR levels were also increased (Figs. 1A and S1A). This observation implied that osimertinib-induced cancer cell death activates the GAS6–MERTK axis in persister cells (RD), but this pathway might be downregulated once tumors acquire resistance. To test this model, we manipulated GAS6 levels in PC9 cells (Del19-EGFR), using either overexpression (OX) or *GAS6* knockout (KO). PCR and immunoblotting confirmed the expected alterations in GAS6 expression (Figs. S1B and S1C). Functionally, GAS6 overexpression enhanced PC9 cell proliferation, whereas GAS6 knockout reduced growth, as measured by crystal violet staining (Fig. 1B). Consistently, MTT assays demonstrated that GAS6-overexpressing cells displayed greater resistance to osimertinib, while KO cells were more sensitive to the drug (Fig. 1C).

**Figure 1:**
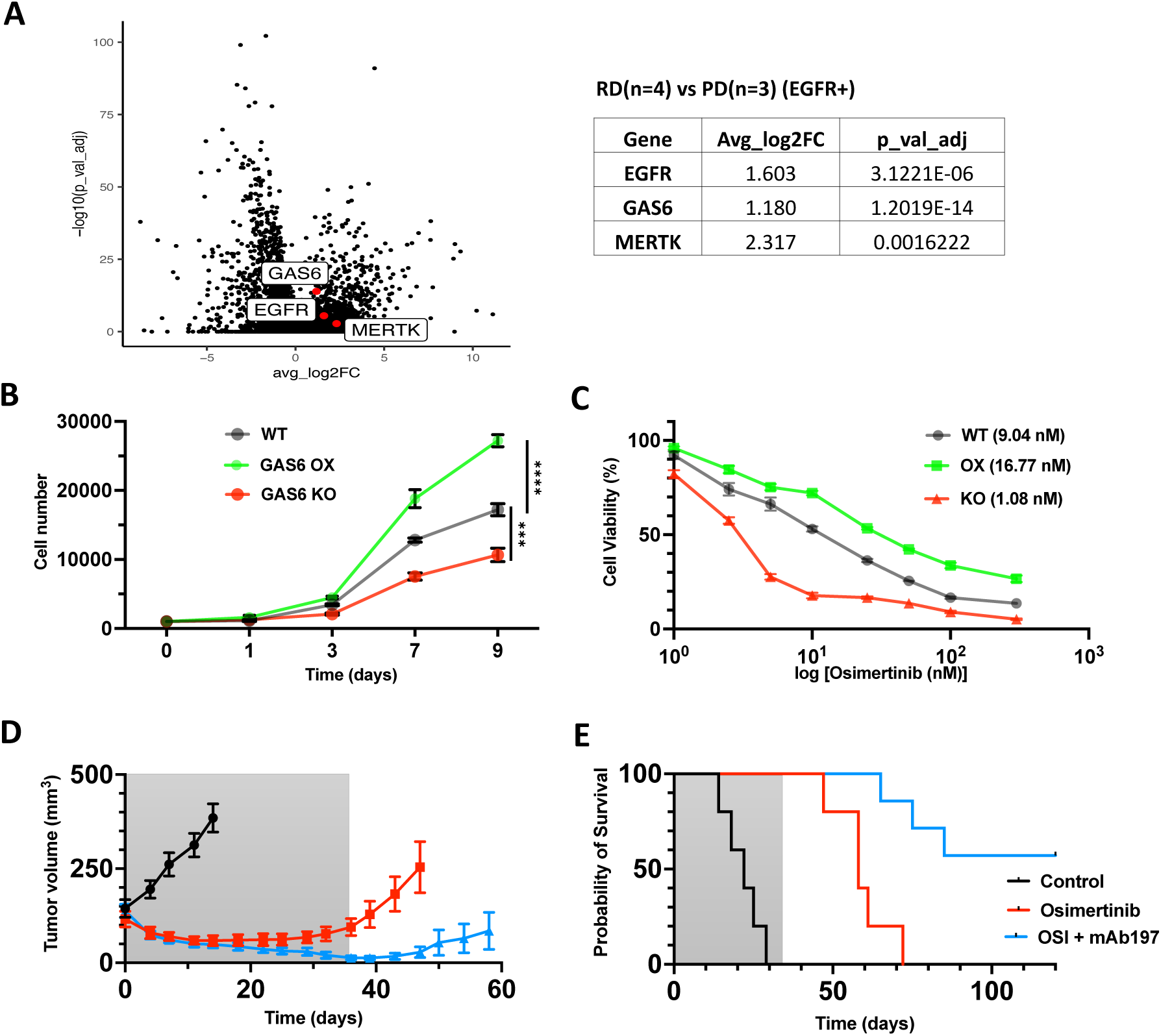
Endogenous GAS6 supports cell survival, and blocking it delays resistance to osimertinib in an animal model. (**A**) Single-cell RNA sequencing data taken from a clinical trial, NCT03433469 (EGFR+ NSCLC patients), were analyzed for differentially expressed genes (DEGs). Patients were treated with osimertinib, and the cohort was segregated into Treatment Naïve (TN), Residual Disease (RD) and Progressive Disease (PD). The volcano plot presents DEGs that differ between patients with RD (4 samples) and PD (3 samples). MERTK, GAS6, and EGFR are highlighted in red. The average log2 of the fold change (FC) and the P-values are presented in a table. (**B**) GAS6 knockout (KO) and GAS6-overexpressing (OX) PC9 cells were generated. Cells (500/well) were seeded in 96-well plates and grown for 9 days. The indicated cells were then fixed and stained with crystal violet to estimate cell number on specific days. ***, p<0.001; ****, p<0.0001. (**C**) The indicated derivatives of PC9 cells (1×10^3^, per well), both genetically modified cells and wildtype cells, were seeded in 96-well plates. Next, cells were treated for 72 hours with different concentrations of osimertinib. Cell viability was determined using a colorimetric assay. (**D** and **E**) PC9 cells (2.5×10^6^) were injected subcutaneously in the flanks of CD1-nu/nu mice. Once tumor became palpable, 5-7 mice were randomized into the indicated 3 groups and treated for five weeks with either osimertinib (10 mg/kg) or/and a mAb against GAS6 (0.2 mg per injection; twice weekly). Average tumor volumes (E) and Kaplan-Meier survival plots (F) are shown. All experiments were repeated at least twice.

Earlier studies showed that GAS6 binds to phosphatidylserine exposed on apoptotic cells ^36^ and activates MERTK- or AXL-expressing cycling persisters, thereby driving mutagenesis and proliferation of TKI-resistant cells ^33^. Based on this mechanism, we hypothesized that blocking GAS6 could suppress osimertinib resistance. To test this, we generated monoclonal antibodies (mAbs) targeting GAS6. Mice were immunized with recombinant human GAS6, and hybridomas were screened for secretion of anti-GAS6 antibodies (Fig. S1D). The lead clone, 197 (an IgG1), was selected and its binding affinity measured by label-free surface plasmon resonance (SPR). Recombinant GAS6 was immobilized, and the antibody was applied at increasing concentrations across the surface. The resulting sensograms (Fig. S1E) revealed an equilibrium dissociation constant of 2.3 nM.

Next, PC9 cells were implanted subcutaneously into immunocompromized CD1-nu/nu mice. Once tumors became palpable, the animals were randomized into three groups (5–7 mice each): untreated controls, mice treated with osimertinib alone (administered orally, once daily) and mice receiving the combination of osimertinib and the anti-GAS6 mAb197 (administered intraperitoneally, twice weekly). Tumor volumes were recorded twice weekly, and body weight was measured once weekly. Treatments were discontinued after five weeks, with no consistent impact observed on body weight, but tumor growth was monitored for an additional four months. Data are presented as average tumor volumes (Fig. 1D), individual tumor growth curves (Fig. S1F), Kaplan–Meier survival analysis (Fig. 1E), and corresponding body weights (Fig. S1G). While osimertinib alone transiently delayed tumor re-growth, its combination with the anti-GAS6 antibody produced markedly stronger and more prolonged effects, leading to complete tumor regression in 4 of 7 mice. Together, these findings highlight GAS6 as a key driver of resistance to a third-generation EGFR inhibitor. They further suggest that therapeutic strategies targeting GAS6, or its receptors, may help inhibit tumor relapse following TKI treatment.

### Endogenous MERTK supports resistance of EGFR-positive NSCLC cells to osimertinib

Both GAS6 and PROS1 activate MERTK and TYRO3, whereas AXL is exclusively activated by GAS6. While the role of AXL in mediating resistance to EGFR inhibitors is well established ^33,38,39,41^, only a limited number of studies have implicated MERTK in progression and relapse of NSCLC ^34,40,42^. To further investigate this possibility, we analyzed Kaplan–Meier survival curves of first progression (FP) in NSCLC patients. The curves shown in Figure 2A track the proportion of patients who remained progression-free at different time points after treatment began (note that a different probe was used in each panel). This data indicates that elevated MERTK transcript levels are associated with earlier disease progression. Based on this clinical correlation, we next examined the functional contribution of MERTK to osimertinib tolerance in EGFR+ NSCLC cells.

**Figure 2:**
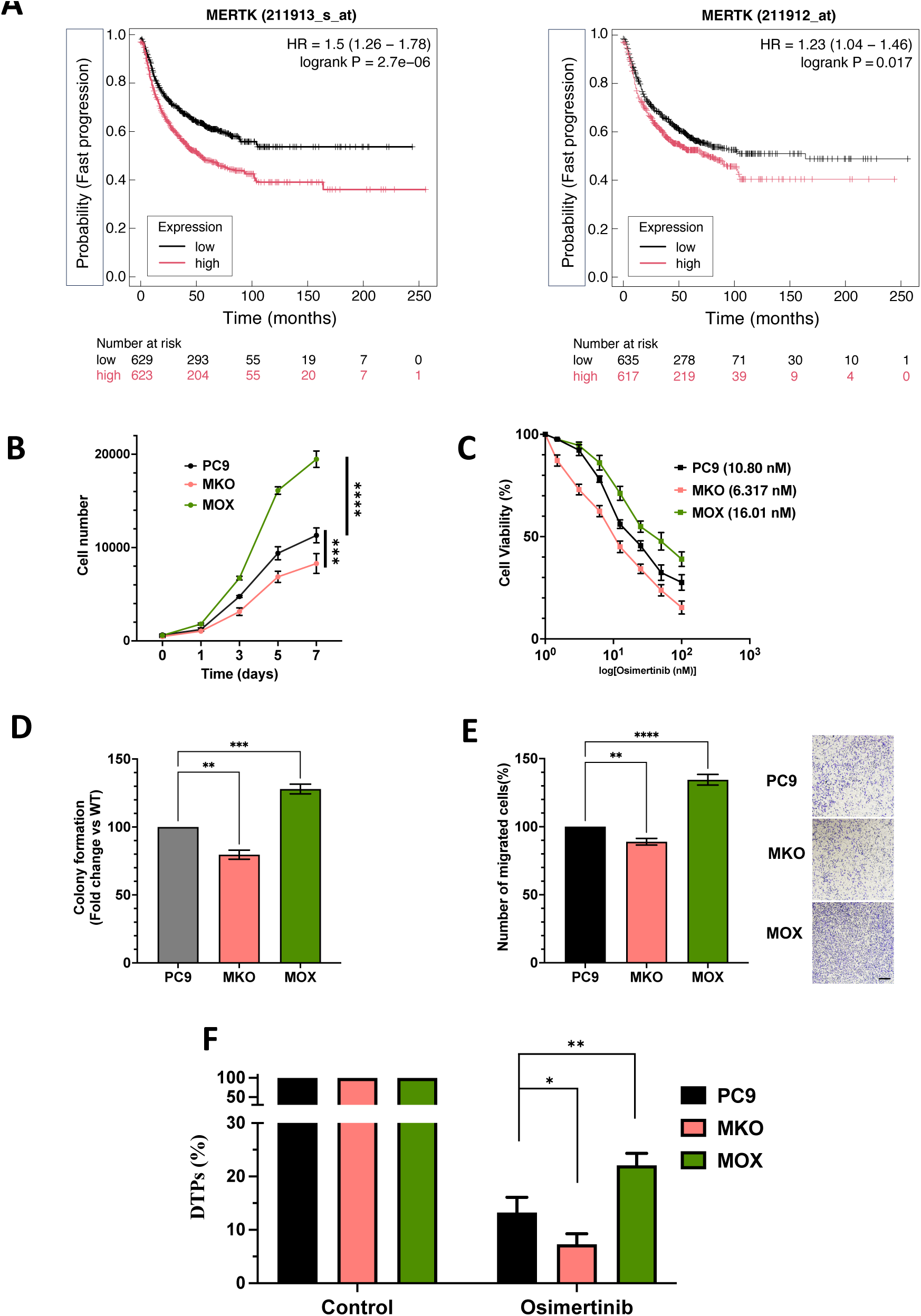
MERTK supports cell survival and drug resistance. (**A**) The KM Plotter tool (https://doi.org/10.1111/bph.16257) was used to obtain Kaplan-Meier survival curves of patients with lung cancer. A dataset of mRNA expression corresponding to 1252 patients was divided into two equal portions according to MERTK levels, high and low, prior to presenting the respective survival curves. Note that each panel presents the results obtained with a different oligonucleotide probe. (**B**) Different PC9 derivatives, as indicated, were seeded in 96-well plates in RPMI medium containing 1% fetal bovine serum. Cell viability was measured using crystal violet staining on consecutive days, and the results were presented in a histogram. The experiment was repeated three times (five technical repeats). ***, p<0.001; ****, p<0.0001. PC9, wildtype cells; MKO, MERTK knockout; MOX, MERTK overexpression. (**C**) Genetically modified and wild-type PC9 cells (3 × 10^3^ per well) were seeded in 96-well plates. Next, cells were treated for 72 hours with different concentrations of osimertinib. Cell viability was determined using the MTT colorimetric assay. The IC50 values were calculated and presented in brackets. (**D**) The indicated PC9 cells (1 x 10^3^ per well) were seeded in 6-well plates and grown for 9 days. Next, cells were fixed and stained with crystal violet. The images of the colonies were scanned and analyzed using ImageJ. **, p<0.01; ***, p<0.001. (**E**) Three different PC9 sublines (6×10^4^ cells per well) were seeded on top of transwell chambers in serum-free medium. The bottom chambers contained media with serum. Cells were incubated for 18 hours. Cells that migrated to the lower face of the transwells were fixed and stained using crystal violet. Next, images were taken under a bright-field microscope. The analysis was performed using ImageJ. Scale bar 2 mm. **, p<0.01; ****, p<0.0001. (**F**) PC9 wild-type and genetically modified cells were seeded in 6-well plates at high confluence. Next, they were treated for 9 days with osimertinib (300 nM). The media and drugs were refreshed once every 3 days. Finally, cells were fixed and stained with crystal violet. Images were taken from 5 non-overlapping fields with a light microscope and analyzed using ImageJ. All experiments were repeated at least twice. *, p<0.05; **, p<0.01.

Using the CRISPR technology, we generated PC9 cells lacking endogenous MERTK (MKO cells) and, conversely, engineered PC9 cells to overexpress MERTK via a plasmid construct (MOX cells). Immunoblot analysis of whole-cell lysates confirmed the anticipated changes in MERTK protein abundance (Fig. S2A). Functional assays demonstrated that MERTK supports proliferation of lung cancer cells: MOX cells showed increased proliferation and expression of Ki67, a proliferation marker (Figs. 2B and S2B). In the next step, we tested the possibility that elevated MERTK expression might enhance the tolerance of EGFR+ lung cancer cells to osimertinib. To evaluate this, MKO, MOX, and control PC9 cells were exposed for 3 days to increasing concentrations of osimertinib, followed by assessment with the MTT assay. As shown in Figure 2C, MERTK overexpression increased the ability of PC9 cells to withstand osimertinib treatment (IC50 increased from 10.8 to 16.0 nM osimertinib) whereas genetic ablation of *MERTK* sensitized the cells to the EGFR inhibitor (IC50 of 6.3nM).

Next, we performed a colony formation assay, which confirmed that *MERTK* ablation diminished long-term proliferative capacity (Figs. 2D and S2C). Moreover, enhanced migration across porous membranes, another hallmark of aggressive cancer cells, was observed in the MERTK-overexpressing PC9 derivative (Fig. 2E). Given that an earlier study identified a reversible drug-tolerant state that precedes the irreversible emergence of TKI-resistant cells ^25^, we also performed a drug-tolerant persister assay. Prior to assying viability, PC9 cells were treated with a relatively high dose of osimertinib for an extended period (9 days). This assay revealed that MOX cells generated more drug-tolerant persister cells compared with MKO and control cells (Figs. 2F and S2D). Taken together, the results shown in Figures 2 and S2 raised the possibility that adaptive MERTK upregulation may endow lung cancer cells with enhanced tolerance to treatment, including TKIs. Specifically, the findings from MKO cells suggested that reducing MERTK abundance could delay the onset of osimertinib resistance.

### Co-targeting MERTK and EGFR using two mAbs delays the onset of resistance to osimertinib, likely due to an EGFR-to-MERTK crosstalk

Unlike TKIs, mAbs can promote endocytosis and degradation of the respective RTKs through their bivalent nature. We therefore hypothesized that anti-MERTK antibodies, partly similar to *MERTK* ablation, would enhance NSCLC sensitivity to TKIs. To generate such antibodies, mice were immunized with a recombinant extracellular domain of MERTK. After several rounds of immunization, blood samples and an immunoprecipitation assay that used PC9 cell extracts confirmed successful antibody production (Fig. S3A). Therefore, splenic B cells were then isolated and fused with a myeloma cell line to create hybridomas. Screening of these hybridomas (Fig. S3B) identified several positive clones, from which one monoclonal line (mAb745) was selected and expanded. Functional testing of mAb745 in the PC9 xenograft model showed that the antibody alone had only weak or no detectable in vivo activity. However, it significantly enhanced the therapeutic effects of both anti-EGFR agents, osimertinib and cetuximab, by delaying tumor relapses (Figs. 3A, 3B and S3C). For instance, none of the osimertinib-only–treated mice survived the study period, whereas 2 of 7 animals receiving mAb745 plus osimertinib survived. Likewise, only 1 of 5 mice treated with cetuximab alone survived, compared to 4 of 7 animals treated with the antibody combination.

**Figure 3:**
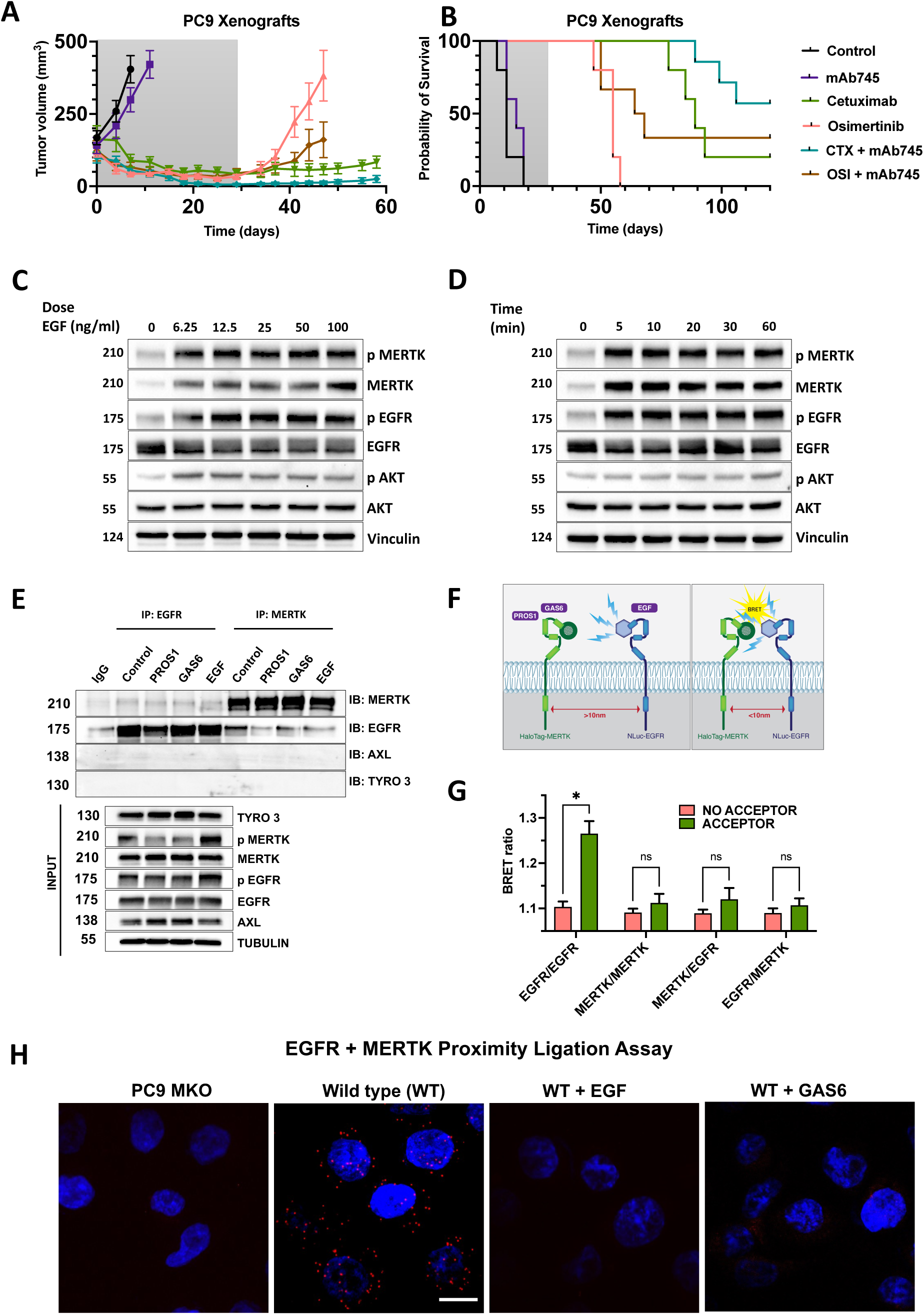
Dual blockade of EGFR and MERTK using antibodies delays tumor relapses, likely due to the ability of EGFR to trans-phosphorylate MERTK within membrane microdomains. (**A** and **B**) CD1-nu/nu mice were subcutaneously implanted with PC9 cells (2.5 × 10⁶). Once tumors became palpable, animals (5–7 per group) were randomized and treated twice weekly for 4 weeks with mAbs (200 µg/injection, intraperitoneally), and/or daily with osimertinib (5 mg/kg; orally). Tumor volumes were measured twice weekly. Tumor volumes and animal survival are reported. The grey-shaded area indicates the treatment period. Data are presented as mean ± SEM for tumor volume (A) and the corresponding Kaplan–Meier survival curves (B). (**C** and **D**) PC9 cells were treated with EGF at increasing concentrations (0, 6.25, 12.5, 25, 50, 100 ng/ml) for 60 min or with 100 ng/ml EGF for varying times (0, 5, 10, 20, 30, 60 min). Cell lysates were immunoblotted with the indicated antibodies, with vinculin as the loading control. (**E**) PC9 cells (2 × 10⁶) were seeded in 10-cm plates and serum-starved overnight. Cells were then treated for 60 minutes with PROS1, EGF, or GAS6 (each at 25 ng/ml). Lysates were prepared and divided for immunoprecipitation using anti-EGFR or anti-MERTK antibodies. Immunoblots were developed as indicated. Note that the input extracts are separately shown (lower panel). Tubulin was used to ensure equal gel loading and immunoglobulin G was used as an immunoprecipitation control. (**F** and **G**) A scheme outlining the BRET assay is shown. HEK293 cells were transfected with NLuc-tagged and Halo-tagged EGFR and MERTK constructs. The following day, cells were treated with the indicated ligands and incubated for 60 minutes at 37°C. The NanoLuc substrate (1:400 final dilution) was then added, followed by a 15-minute incubation in the dark. Luminescence was measured using a PHERAstar FSX plate reader, a 475 ± 15 nm bandpass filter (donor) and 665/535 ± 15 nm bandpass filter (acceptor). Raw BRET ratios were calculated by subtracting fluorescence emissions from luminescence emissions as the 665/475 nm signal and baseline-corrected using the ‘no acceptor’ control. *, p<0.05.ns, non-significant. (**H**) PC9 cells (10^4^) were seeded on coverslips and allowed to adhere overnight. The next day, cells were serum starved for 6 hours and treated with either EGF (100 ng/ml) or GAS6 (100 ng/ml) in serum-free RPMI, for 1 hour. Thereafter, cells were fixed with 4% PFA, permeabilized and then stained with a rabbit anti-MERTK antibody and with a murine anti-EGFR antibody. Subsequently, a proximity ligation assay was performed using anti-Rabbit Plus and anti-Mouse Minus. The amplified product of the PLA was labeled using a red dye, and nucleai were stained with DAPI. Images were obtained using a Zeiss Spinning Disk confocal microscope, and the analysis was performed using ImageJ. MERTK knockout PC9 cells were used as a negative control. Scale bar 10 μm. All experiments were repeated at least twice.

The ability of mAb745 to enhance the efficacy of two highly different anti-EGFR agents suggested the existence of an EGFR–MERTK crosstalk, potentially critical for the development of drug resistance. Two observations supported this notion: (i) stimulation of PC9 cells with EGF induced rapid phosphorylation not only of EGFR but also of MERTK (Figs. 3C and 3D), and (ii) co-immunoprecipitation with anti-EGFR antibodies retrieved small amounts of MERTK (Fig. 3E). Conversely, immunoprecipitation with anti-MERTK antibodies pulled down trace levels of EGFR, but pretreatment with EGF, GAS6, or PROS1 either reduced or did not alter these receptor-receptor associations. To further probe these receptor asociations, we employed nanoBRET assays ^43^, which monitor protein—protein interactions in live cells via bioluminescence resonance energy trasfer and have been used by us to characterize the molecular pharmacology of EGFR ^44^. As acceptor, the membrane impermeable HaloTag AlexaFluor-488 substrate was used. The fluorophore labels were attached to the N-terminus of either HaloTagMERTK or HaloTagEGFR. When donor and acceptor are in close proximity (0-10 nm), NanoLuc emission excites the fluorophore on the HaloTag sequence, which then emits light at a longer wavelength (schematize in Fig. 3F). Consistent with spontaneous EGFR homodimerization, we observed significant BRET signals when NanoLuc and HaloTag were separately fused to EGFR and co-expressed (Fig. 3G). In contrast, no evidence was obtained for MERTK homodimers or for MERTK-EGFR heterodimerization, even after stimulation with GAS6, PRO1 or EGF (Fig. S3D).

Assuming that inter-receptor distances exceed the <10 nm distance requirement for efficient BRET, we used the proximity ligation assay (PLA), which relies on pairs of antibodies and detects interactions within an effective range of ∼10–40 nm. Gratifyingly, PLA revealed numerous EGFR–MERTK clusters in PC9 cells, whereas no signal was observed in MERTK-knockout cells, confirming assay specificity (Fig. 3H). Impressively, pretreating naïve PC9 cells with either GAS6 or EGF almost completely abolished the PLA signal, consistent with the co-immunoprecipitation assays and a previously described mechanism that concentrates distinct unstimulated RTKs within caveolae ^45^. According to this and additional reports, following ligand binding these receptors rapidly exit caveolae in a process coupled to receptor auto-phosphorylation and Src kinase activation. Taken together, these findings suggest that the in vivo observed ability of an anti-MERTK antibody to enhance the efficacy of anti-EGFR agents (i.e., osimertinib and cetuximab) is mediated by inhibition of a caveolae-localized stimulatory crosstalk. This physical interaction enables spontaneous transphosphorylation of MERTK by EGFR, which might adaptively contribute to drug resistance.

### A bispecific MERTK/EGFR antibody enhances endocytosis and degradation of both MERTK and EGFR

bsAbs expand the therapeutic arsenal beyond monoclonals by dual receptor blockade and combining tumor targeting with immune activation ^46^. Compared with conventional mAbs, bsAbs can block compensatory signaling pathways, promote receptor internalization and degradation and increase specificity by requiring co-recognition of tumor-associated antigens. For the design of the bispecific anti-MERTK/EGFR antibody (termed Bis3; see a schematic diagram in Fig. 4A and aminoacid sequences of all four chaines umder Methods), we replaced the murine Fc region of mAb745 and combined the recombinant forms of mAb745 and cetuximab (an anti-EGFR mAb), both carrying a human Fc domain. To prevent mispairing of the light and heavy chains during co-expression of the two antigen-binding arms, the EGFR-binding arm of cetuximab was reformatted as a CrossMab with a CH1–CL domain swap ^47^. Additionally, heterodimerization of the two heavy chains was enforced through the following CH3 knobs-into-holes mutations ^48^: Y349C/T366S/L368A/Y407V in the EGFR arm and S354C/T366W in the MERTK arm. Binding affinities of Bis3 and its parent molecules towards recombinant forms of MERTK and EGFR were assessed by surface plasmon resonance (SPR; Figs. S4A–S4D). The calculated dissociation constants (Kd), derived from the ratio of the dissociation rate constant (koff) to the association rate constant (kon), are summarized in Figure 4A. As shown, Bis3 displayed an approximately fourfold lower affinity for MERTK relative to the parental recombinant antibody, while its affinity for EGFR was substantially reduced relative to cetuximab. This decrease in binding strength of both antigen-binding arms is not uncommon and attributable to altered protein folding. Importantly, the results of a sandwich ELISA test that used an immobilized MERTK were consistent with simultaneous engangement of both MERTK and EGFR molecules by Bis3 (Fig. 4B). In addition, we confirmed that Bis3 was able to precipitate both EGFR and MERTK, as expected from a bivalent molecule having dual specificity (Fig. 4C).

**Figure 4:**
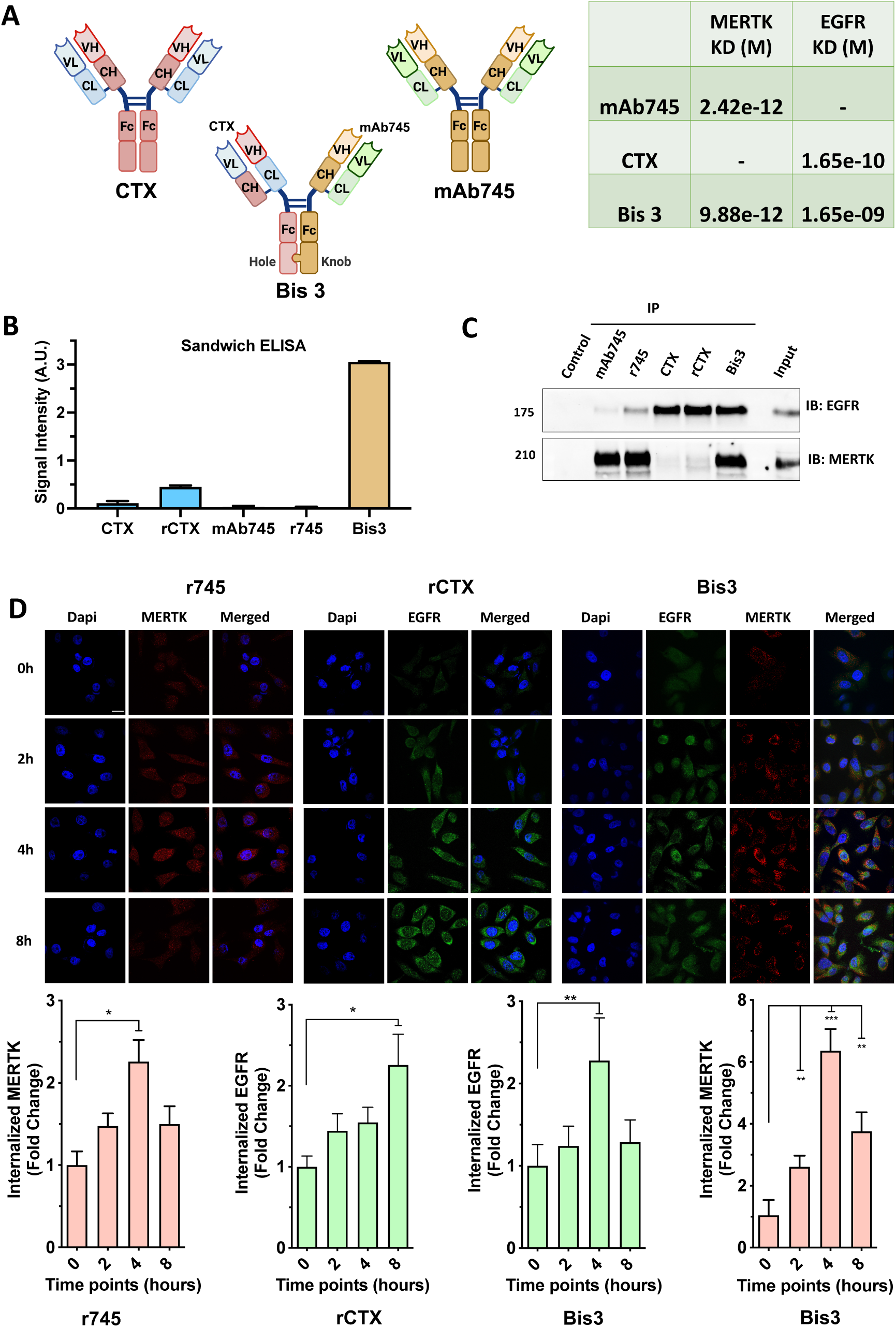
Design and activity of the anti-MERTK/EGFR bispecific antibody (Bis3). (**A**) Schematic representation of the bispecific antibody, Bis3, and the parental antibodies, anti-MERTK mAb745 and cetuximab, a chimeric human/mouse anti-EGFR antibody. The EGFR-binding arm was formatted as a CrossMab (CH1–CL swap) to ensure correct heavy–light chain pairing. Additionally, heterodimerization of the heavy chains was achieved through inserting the following mutations in the CH3 domain: Y349C, T366S, L368A and Y407V in the EGFR arm and two mutations, S354C and T366W, in the MERTK arm. V denotes an antibody variable domain and C marks a constant antibody domain. Likewise, H denotes a heavy chain and L denotes the light chain. The table lists parameters obtained using Surface Plasmon Resonance (SPR), which determined binding affinities (dissociation constant, Kd) of the parental antibodies, as well as the Kd of Bis3, towards recombinant soluble forms of human MERTK and human EGFR. CTX, cetuximab. This scheme was made using BioRender. (**B**) Ninety-six-well plates were coated overnight with a soluble ectodomain of MERTK. The following day, the plates were blocked with albumin-containing saline, and thereafter they were incubated with the indicated antibodies (20 μg/ml). After washing, biotinylated EGFR’s ectodomain was added, followed by avidin-HRP. The signal was measured using a microplate reader. Note that both native and engineered (recombinant) versions of cetuximab (rCTX) and mAb745 (r745) were used. (**C**) PC9 cell lysates were subjected to immunoprecipitation assays that used Bis3, anti-EGFR and anti-MERTK antibodies. The immunoblots were probed with the indicated antibodies. Recombinant forms of mAb745 and cetuximab were used, alongside the native forms. (**D**) PC9 cells (5×10^4^) were seeded on a coverslip. The next day, cells were treated for various time intervals with the following antibodies: CTX (cetuximab; 10 μg/ml), mAb745 (10 μg/ml) and Bis3 (20 µg/ml). Next, cells were fixed (with 4% PFA), permeabilized and stained with anti-MERTK and anti-EGFR antibodies, which were followed by fluorescently tagged secondary antibodies. DAPI was used to visualize nuclei. Images were taken using a spinning disk confocal microscope and a series of Z-stacks were taken with a z-step of 1 μm. To quantify the intracellular levels of the indicated receptors the middle section of the cells was selected from the Z-stack and for each channel the average signal in the cytosol of the cells was calculated (see histograms at the bottom of the figure). The obtained signal intensity was used as threshold value to remove background cytosolic signals. The spots were counted using the function “analyze particles” of ImageJ. The signal corresponding to receptor internalization at t=0 was used for normalization. *, p<0.05; **, p<0.01; ***, p<0.001. Scale bar, 15 μm. All experiments were repeated at least twice.

Many mAbs directed against cell-surface receptors can trigger receptor internalization (endocytosis) ^49^. Clinically, this attribute might be important: internalization can attenuate receptor signaling, but excessive uptake may reduce receptor availability for potential antibody-dependernt cellular cytotoxicity (ADCC). To address Bis3-induced receptor endocytosis, we exposed PC9 cells to increasing concentrations of either mAb745, cetuximab or Bis3, and monitored MERTK and EGFR abundance by immunoblotting. This analysis validated mAb745- and cetuximab-induced degradation of MERTK and EGFR, respectively (Fig. S4E). Furthermore, both MERTK and EGFR were downregulated following Bis3 treatment. To examine this mechanism more directly, we employed immunofluorescence. Cells were treated with Bis3 or with the two parental antibodies, fixed and probed using the respective polyclonal antibodies. Critically, imaging was performed using a spinning disk confocal microscope that was focused beneath the plasma membrane to capture internalized receptors en route to lysosomal degradation. As shown in Figure 4D and quantified in the accompanying histograms, both MERTK and EGFR underwent internalization within 2-4 hours of treatment with the respective antibody, and as anticipated, Bis3 promoted internalization of both receptors. In summary, Bis3 retained significant binding affinities for both EGFR and MERTK, and was capable of driving their internalization and intracellular degradation. These findings suggest that Bis3 may simultaneously inhibit the primary signaling pathway mediated by EGFR and the adaptive pathway mediated by MERTK, thereby potentially inhibiting the emergence of resistance to EGFR kinase inhibitors.

### When tetsted in 3D spheroids, a combination of TKIs (osimertinib plus MRX-2843) is more effective than Bis3 or the corresponding mAb combination

Multi-kinase inhibitors (e.g., sorafenib), or a combination of kinase inhibitors each specific to a distinct target, may offer advantages over bsAbs or dual combinations of mAbs. As an initial step toward comparing the efficacies of a mAb pair and a pair of two TKIs we examined the ability of osimertinib to cooperate with MRX-2843, a small-molecule MERTK inhibitor currently under clinical development. Two models are commonly used to study cooperative effects of small molecule drugs: the Bliss and the Loewe models ^50^. Practically, these models consider the dose-response curves of individual drugs and makes visual representations of each pair of drugs based on data corresponding to multi-dose combination responses. Accordingly, we treated PC9 and H1975 cells (EGFR L858R and T790M) for 72 hours with increasing concentrations of osimertinib and MRX-2843, and assessed cell viability using crystal violet staining. The results were used for the calculation of drug-drug interactions (synergy scores) using the Loewe model, as shown in Figure S5A. Interestingly, for both cell lines, relatively low concentrations of osimertinib achieved high synergy only when MRX-2843 was present at relatively high concentrations.

Hence, we performed colony formation assays with H1975 cells (expressing the resistance-associated EGFR-T790M mutation and L858R) while applying relatively low and high concentrations of osimertinib and MRX-2843, respectively (Figs. 5A). Cells were treated for nine days with osimertinib, MRX-2843, Bis3 and the parental anti-EGFR and anti-MERTK antibodies. Quantification of stained colonies revealed that the most potent regimens were the triplet combination MRX-2843 + osimertinib + cetuximab and the doublet Bis3 plus osimertinib (Fig. 5A). Similar to these findings, the same drug combinations emerged as the strongest inhibitors in a drug-tolerant persister assay (Fig. 5B). As aforementioned, this assay, performed with both H1975 and PC9 cells, measures the capacity of treatments to suppress the reversible formation of persister cells—precursors to stable resistance ^25^. A third assay assessing short-term viability after three days of drug exposure (Fig. S5B) further confirmed that dual blockade of EGFR (with osimertinib + cetuximab) and MERTK (with Bis3, mAb745 or MRX-2843) was required to achieve maximal suppression of cell survival in both in vitro models we examined.

**Figure 5:**
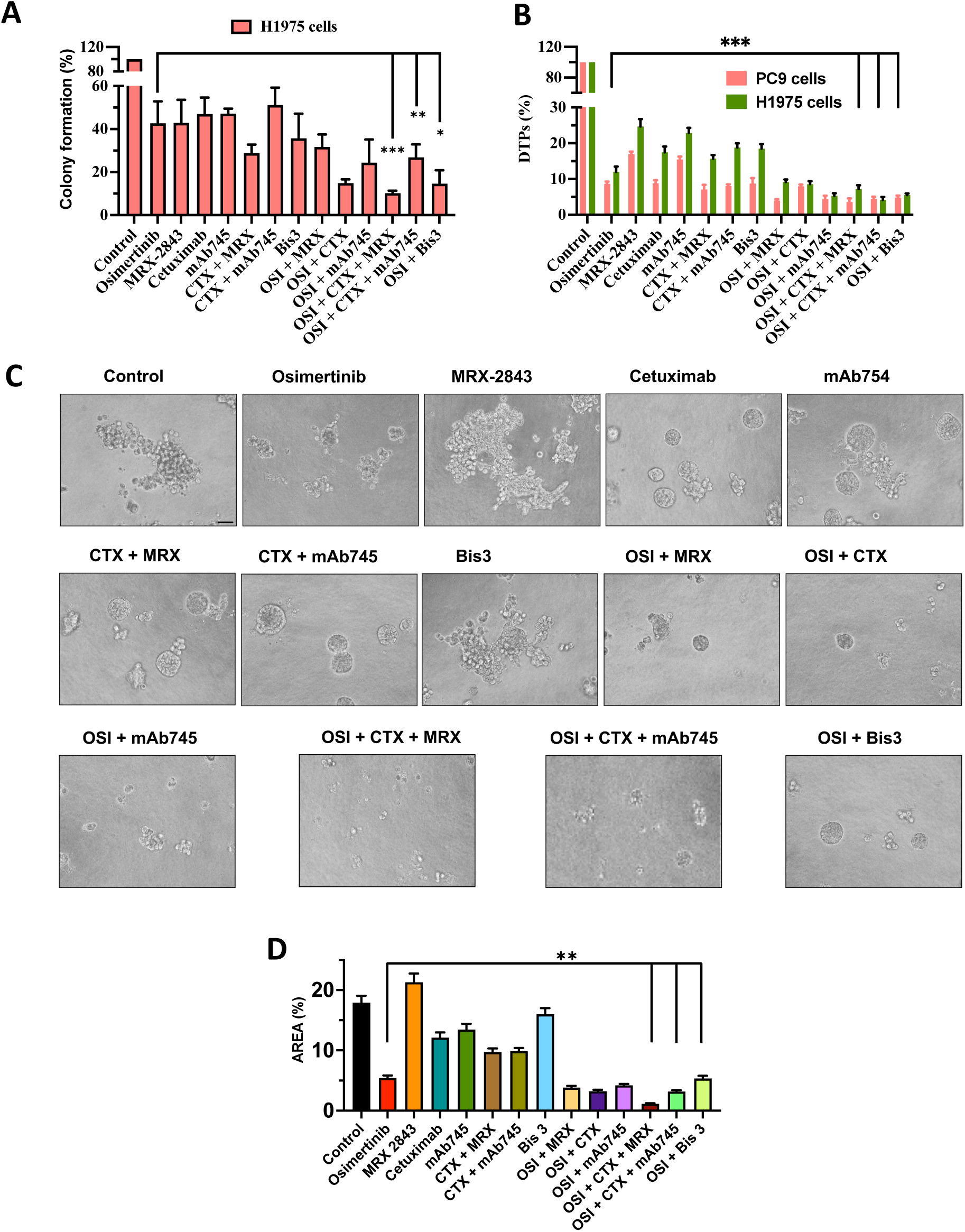
Simultaneous inhibition of MERTK and EGFR effectively suppresses NSCLC cells harboring exon-19 and exon-21 EGFR mutations. (**A**) H1975 cells (EGFR-L858R in exon-21 and the “gatekeeper” mutation T790M) were seeded in 6-well plates (1,000 cells/well). The next day, cells were treated for 9 days with anti-EGFR and anti-MERTK mAbs (10 µg/ml), Bis3 (20 µg/ml), osimertinib (10 nM) or MRX-2843 (300 nM). The drugs were replenished once every 72 hours. Next, cells were fixed and stained with crystal violet. Images were captured using a bright-field microscope. Colonies were quantified (CFU, colony formation units) using ImageJ. The experiment was repeated thrice. (**B**) PC9 (EGFR exon-19 deletion) and H1975 cells were seeded and on the next day cells were treated for 9 days as in A. Next, they were fixed and stained with crystal violet. Images of surviving cells were captured using a bright-field microscope. Drug tolerant persister (DTP) cells were quantified using ImageJ. (**C** and **D**) PC9 cells (1×10^3^) embedded in 5% BME were seeded in BME pre-coated 96-well plates. The wells contained vehicle control (CTR), mAbs (10 μg/ml), osimertinib (10 nM), MRX-2843 (300 nM), Bis3 (20 μg/ml) and the indicated drug combinations. All experiments were stopped after 7 days. Images of the formed spheroids were captured using an OpTech IB4 microscope (bright field; 20x magnification). Percentages of well covered areas were assessed using ImageJ. Significance was calculated using one-way ANOVA, Dunnett’s and Tukey’s multiple comparison tests (**, p<0.01). The histograms represent the average of two independent experiments performed in duplicates. All experiments were repeated at least twice. Scale bar, 50 μm.

Unlike 2D monolayers, tumor resistance to anti-cancer drugs frequently associates with diffusion barriers and tumor–matrix interactions, both of which are more accurately preserved in 3D spheroids. These multicellular aggregates recapitulate key features of tumors—including architecture, cell–cell contacts, and extracellular matrix organization—thereby providing a physiologically relevant context for investigating mechanisms of drug resistance ^51^. To exploit this model, we established 3D cultures of PC9 cells embedded in basement membrane extract (BME), which was enriched in matrix proteins, such as collagens and laminin, and treated them for seven days with various drug combinations. Unexpectedly, the triplet regimen of osimertinib + cetuximab + MRX-2843 produced significantly stronger inhibition than combinations in which MERTK was blocked with antibodies (Figs. 5C and 5D).

To extend these observations to an osimertinib-resistant model, we utilized PC9-AZDR cells, a derivative line that had evolved resistance to osimertinib following prolonged exposure to the drug (>3 months). As expected, spheroids generated from PC9-AZDR cells were largely unresponsive to osimertinib-containing combinations (Figs. S5C and S5D). However, they remained markedly sensitive to the triplet regimen of osimertinib + cetuximab + MRX-2843. Overall, our in vitro assays demonstrated that dual blockade of EGFR and MERTK—whether through kinase inhibitors or a bispecific antibody—is required for optimal suppression of NSCLC cell survival. Importantly, this requirement was consistently observed in both 2D and 3D spheroid models. Within this context, MERTK inhibition via a small-molecule kinase inhibitor was more effective than antibody-based approaches, likely reflecting the influence of spheroid architecture and diffusion gradients unique to each drug.

### Combined Bis3 and osimertinib treatment durably suppresses exon-19 and exon-21 xenograft models

To assess whether dual inhibition of EGFR and MERTK can control tumor growth in an animal model, we generated xenografts by subcutaneously implanting PC9 (exon-19; Figs. 6A and 6B) and H1975 cells (exon-21; Figs. 6C and 6D) into the flanks of immunocompromized mice. Once tumors became palpable, animals were randomized into treatment groups. Small-molecule inhibitors (osimertinib and MRX-2843) were administered orally once daily, while antibodies (Bis3, cetuximab and mAb745) were delivered intraperitoneally twice weekly. Treatments were discontinued after four weeks, but tumor growth and body weight were monitored for an additional four months. Outcomes are presented as Kaplan–Meier survival plots (Figs. 6A–D), individual tumor growth curves (Figs. S6A and S6C), and body weight trajectories (Figs. S6B and S6D). No consistent weight loss—used as a surrogate for systemic toxicity— was observed across treatment groups. In contrast, tumor burden and survival varied substantially. As expected, osimertinib monotherapy initially reduced tumor size, but relapse occurred soon after treatment cessation. Combination regimens yielded stronger and more durable effects: all osimertinib-based doublets extended survival compared with osimertinib alone, with osimertinib + cetuximab being the most effective pair.

**Figure 6:**
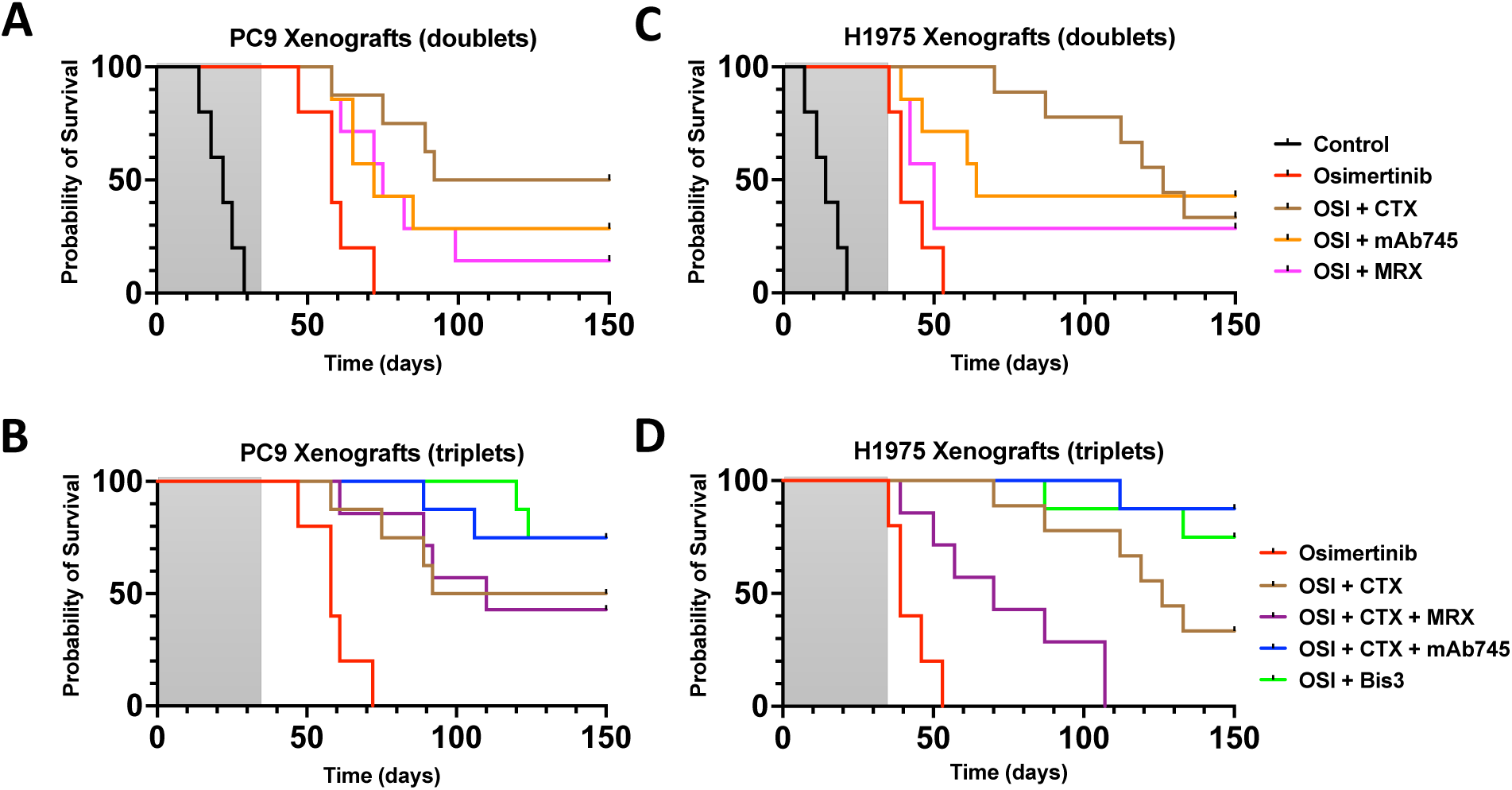
In an animal model, the efficacy of the combination Bis3+osimertinib surpasses that of osimertinib+cetuximab+MRX-2843. (**A** and **B**) PC9 cells (2.5×10^6^ per animal) were injected subcutaneously in the flanks of CD1-nu/nu mice. Once tumors became palpable, 5-8 mice were randomized and treated for five weeks with either mAbs (100 μg/injection) or Bis3 (200 μg/injection), twice a week. In addition, where indicated, we used osimertinib (10 mg/kg) or MRX-2843 (20 mg/kg), which were orally administered on a daily basis. Tumor volumes were monitored twice a week and animal body weight was determined once a week. The grey-marked area denotes treatment duration. (**C** and **D**) The H1975 xenograft model was employed essentially as described for the PC9 model shown in panels A and B.

Notably, unlike the in vitro results, the triplet osimertinib + cetuximab + mAb745 was more effective than the MRX-2843–containing triplet (osimertinib+cetuximab+MRX-2843). Furthermore, the Bis3 + osimertinib doublet achieved efficacy comparable to that of the osimertinib + cetuximab + mAb745 triplet across both xenograft models. In summary, dual inhibition of EGFR and MERTK significantly suppressed tumor growth in both PC9 and H1975 xenografts. The bispecific antibody Bis3, in combination with osimertinib, reproduced the strong and durable tumor control observed with the most potent triplet regimen. Presumably, the superior performance of the anti-MERTK antibody compared with the kinase inhibitor in triplet combinations reflects its slower systemic clearance and/or the unique capacity of antibodies, unlike TKIs, to recruit anti-tumor immune effector cells (note that we used an immunocompromized mouse model that maintains most myeloid lineages).

### The bispecific EGFR/MERTK antibody prolongs remission in patient-derived xenografts harboring EGFR mutations in either exon-19 or exon-20

Patient-derived xenograft (PDX) models maintain key molecular characteristics of the original tumors, including intratumoral heterogeneity, and therefore more accurately reflect drug resistance mechanisms than traditional cell line xenografts ^52^. To test Bis3 in a PDX model, we selected the TM00193 PDX model (E746_A750 Del19-EGFR, from The Jackson Laboratory) and subcutaneously implanted fragments in immunocompromized mice. Once tumors reached ∼350 mm³, animals were randomized into treatment groups and received osimertinib, recombinant mAbs or Bis3 for a three-week period. Tumor growth and animal survival were monitored for up to 120 additional days (Figs. 7A, 7B and S7A). The data showed that Bis3 worked synergistically with osimertinib, preventing the typical rapid relapse observed before and after drug withdrawal. Moreover, consistent with the cell line xenograft studies, Bis3 produced efficacy comparable to the triplet combination (osimertinib + cetuximab + mAb745).

**Figure 7:**
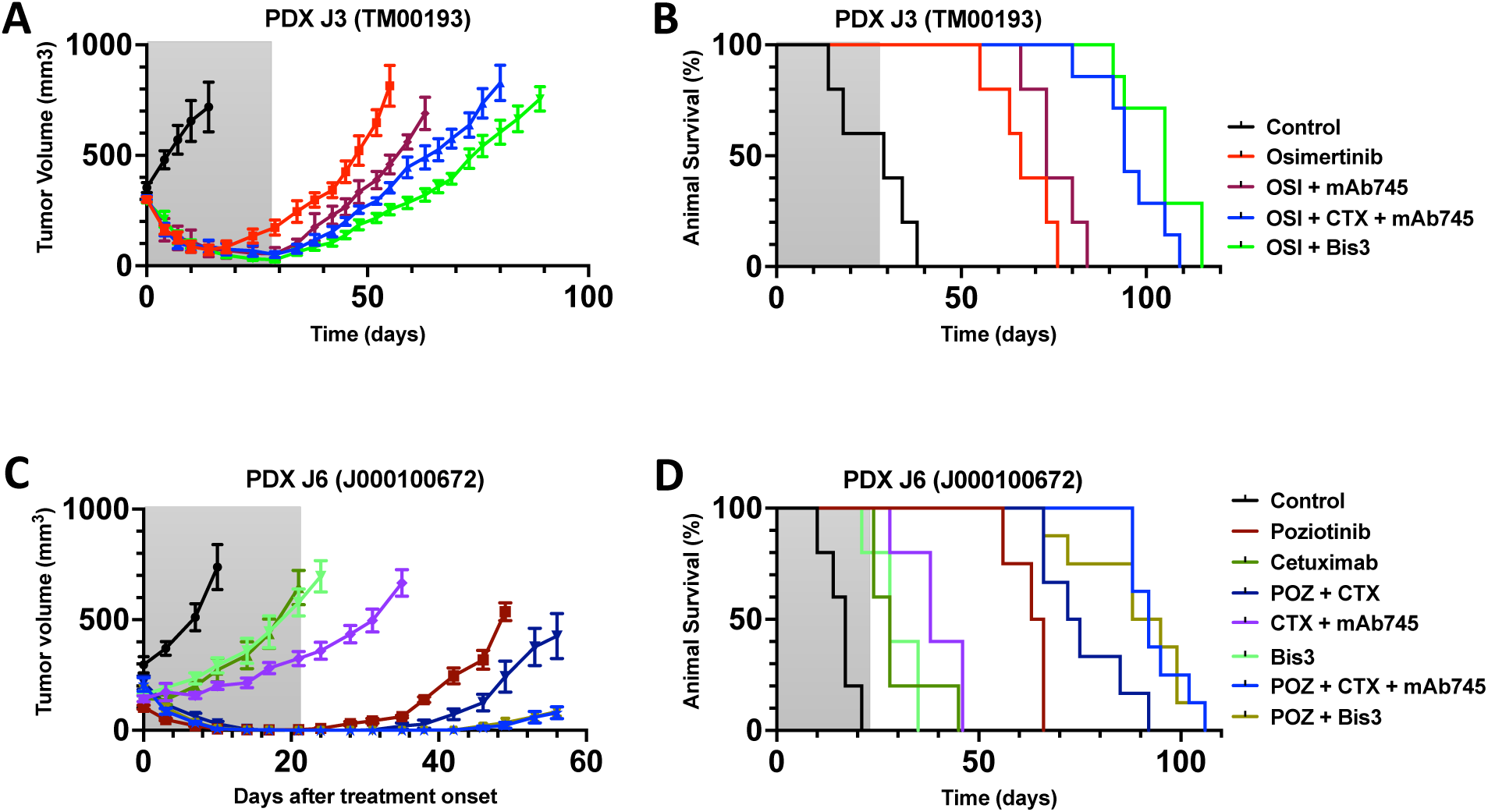
The EGFR–MERTK bispecific antibody enhances the efficacy of TKIs by delaying relapse in PDX models harboring EGFR exon-19 and exon-20 mutations. (**A**) The patient-derived xenograft TM00193 (E746_A750 Del19-EGFR) was engrafted subcutaneously in NSG mice. When tumors became palpable (around 350 mm^3^), mice were randomized in groups of 5-7 animals and treated for four weeks with osimertinib (10 mg/kg/day; orally) and the indicated monoclonal antibodies (100 micrograms/mouse/injection), or with Bis3 (200 micrograms/mouse/injection). Tumor volume and animal body weight were monitored for the next 90 days. Data shown are means ± SEM. (**B**) Shown are Kaplan-Meier survival curves corresponding to the experimental arms presented in A. (**C** and **D**) NSG mice were implanted with tumor fragments derived from a lung cancer PDX model, PDXJ6 (from The Jackson Laboratory, J000100672) harboring the exon-20 mutations S768_V769insSVD. Once tumors reached approximately 300 mm³, mice were randomized into groups of 5-7 animals per group. They were later treated for 3 weeks (shaded areas), twice a week, with the indicated mAbs (100 μg/injection) or with Bis3 (200 μg/injection). Alternatively, mice were treated with an exon-20 specific TKI, poziotinib (5 mg/kg), which was orally administered on a daily basis. Tumor growth and body weight were monitored twice per week. Shown are the average tumor volumes (C) and animal survival times per group (D). Data shown are means ± SEM.

Unlike the exon-19 deletions and the L858R mutation in exon-21, which represent the majority of EGFR mutations in NSCLC, exon-20 insertions are relatively rare and confer a unique mechanism of kinase activation ^53^. These structural attributes translate to lack of response to clinically approved EGFR inhibitors. Still, several compounds that selectively inhibit EGFR exon-20 insertions have recently been reported but only one, sunvozertinib ^54^, has been approved so far. Another drug, poziotinib, showed activity in clinical trials but had dose-limiting toxicities. Importantly, amivantamab, a bispecific EGFR/MET mAb, received approval in combination with chemotherapy for patients with EGFR exon-20 insertions, whose disease had progressed after chemotherapy ^55^.

Since all EGFR mutants retain a highly conserved extracellular domain, we examined whether Bis3 could also be effective against tumors harboring the difficult-to-treat exon-20 mutations. To this end, we used an aggressive PDX model from The Jackson Laboratory (J000100672), established from a patient with the S768_V769insSVD mutation. Tumor fragments from this model were engrafted into mice and treated with different antibody regimens, including Bis3, either alone or in combination with an exon-20–selective TKI, poziotinib. Because prolonged, high-dose poziotinib has been linked to notable toxicity in patients, we administered a relatively low dose for a limited period to reduce the commonly observed delayed side effects ^56^. As shown in Figures 7C, 7D, and S7B, poziotinib alone only partially delayed relapse of the exon-20 xenografts, and neither Bis3 nor the respective antibody combination (mAb745 plus cetuximab) blocked the aggressive growth of this model. However, both antibody treatments markedly potentiated the effect of poziotinib, even under low-dose and short-term conditions. Importantly, in this exon-20 model, neither poziotinib-nor osimertinib-based combinations consistently affected body weights (Fig. S7C). Overall, the findings made using this challenging PDX model indicate that Bis3, in similarity to the combination of its parental antibodies, can significantly delay resistance to poziotinib, supporting the possibility that newer, less toxic exon-20 TKIs may provide prolonged clinical benefit—particularly when given continuously rather than in intermittent, drug-holiday regimens similar to our experimental protocol.

In summary, this study was inspired by the clinical observation that the GAS6–MERTK pathway becomes activated in patients with EGFR+ NSCLC following TKI treatment. GAS6 detects apoptosis and signals through its receptor, MERTK, initiating survival pathways that likely promote adaptive mutagenesis and resistance. To counteract this, we developed three therapeutic antibodies: anti-GAS6, anti-MERTK and a bispecific EGFR/MERTK antibody (Bis3). Blocking either GAS6 or MERTK, together with EGFR inhibition, significantly delayed the onset of resistance, not only in tumors driven by exon-19 deletions or the L858R-EGFR mutation (exon-21), but also in a model of NSCLC harboring the clinically challenging exon-20 mutations. Interestingly, although a combination of MERTK- and EGFR-selective TKIs showed higher efficacy in vitro, Bis3 and the the respective pairwise combination of anti-MERTK and anti-EGFR mAbs achieved superior activity in multiple animal models, suggesting an additional contribution from either endocytosis or immune-mediated mechanisms. Collectively, these findings underscore the translational promise of Bis3 and the therapeutic value of targeting the GAS6–MERTK axis to overcome resistance to EGFR kinase inhibitors.

## Discussion

MERTK has been repeatedly implicated in resistance to diverse anticancer therapies. For instance, this receptor activates pro-survival signaling that diminishes the effectiveness of chemotherapy, whereas its inhibition sensitizes tumor cells to cytotoxic agents ^57^. MERTK also contributes to an immunosuppressive tumor microenvironment that reduces the efficacy of radiotherapy ^58^. Moreover, resistance to targeted kinase inhibitors—such as EGFR or AXL-specific TKIs—can involve MERTK, which functions as a bypass node that reactivates downstream survival pathways ^59^. One key to understanding the roles played by MERTK and AXL in drug resistance is their rather unique ability to mediate efferocytosis, the process by which apoptotic cells are recognized and engulfed by macrophages and dendritic cells ^60^. The ligands of TAM family receptors, GAS6 and Protein S, serve as bridging molecules by binding phosphatidylserine exposed on dying cells while simultaneously engaging MERTK and other TAM receptors on phagocytes, thereby driving apoptotic cell clearance. Our study was guided by clinical observations: both GAS6 and MERTK—the most critical mediators of efferocytosis—were found to be upregulated in tumors of lung cancer patients shortly after initiation of TKI treatment. We therefore hypothesized that extensive drug-induced apoptosis, followed by efferocytosis, may activate MERTK-mediated bypass signaling pathways. Hence, we assumed, blocking either GAS6 or MERTK might delay the onset of resistance to osimertinib.

To test this model, we developed three antibodies: two mAbs—one targeting GAS6 and the other MERTK—and a bispecific antibody, Bis3, directed against both MERTK and EGFR. Bispecific antibodies (BsAbs) like Bis3 offer several advantages over single-target mAbs. By simultaneously engaging two antigens, they can enhance efficacy, improve safety, and simplify treatment regimens ^61^. Critically, by blocking multiple pathways at once, BsAbs can counteract tumor escape mechanisms and prevent the development of resistance that often limits the efficacy of single-target mAbs. This is especially relevant in the context of EGFR-specific TKIs, where adaptive activation of bypass receptors—including HER2, HER3, AXL, MET, MERTK, and IGF1R—is a common event. Inhibiting individual bypass pathways has been shown to partially delay relapse ^33,40,62–66^. A clinically validated example of this approach is amivantamab, a bispecific IgG1 antibody directed against EGFR and MET. Unlike TKIs, amivantamab not only drives receptor internalization and degradation but also enhances antibody-dependent cellular cytotoxicity (ADCC) by engaging NK cells and macrophages ^67^. Similarly, Bis3 promotes degradation of both EGFR and MERTK and, in our preclinical models, markedly delays the onset of resistance to TKIs, including an exon-20–selective inhibitor. Bis3 also incorporates a human Fc domain, enabling the recruitment of immune effector functions. Collectively, these features support Bis3 as a compelling candidate for clinical translation. This may be particularly relevant for patients with the difficult-to-treat EGFR exon-20 insertions. When identified at diagnosis, such patients may receive first-line therapy with amivantamab plus chemotherapy. Upon disease progression, amivantamab monotherapy or TKIs are typically considered. However, using an exon-20 PDX model, we observed strong synergy between Bis3 and an exon-20–selective TKI, pointing to an additional therapeutic strategy.

As expected, both anti-MERTK and anti-GAS6 antibodies delayed tumor relapse in animal models, underscoring the critical role of the GAS6–MERTK axis in resistance to TKIs. However, blocking GAS6 or inhibiting MERTK alone proved insufficient; maximal suppression of resistance required concurrent EGFR inhibition through both a TKI (osimertinib) and an antibody (cetuximab). These findings are consistent with the ability of the GAS6–MERTK pathway to provide an alternative survival route when EGFR signaling is fully suppressed. Notably, while a MERTK kinase inhibitor demonstrated robust activity in 3D in vitro systems, its efficacy was markedly reduced in the in vivo models. By contrast, the antibodies we evaluated, including the bispecific Bis3, showed superior performance in vivo, likely due to their capacity to engage residual immune effector cells—particularly myeloid-derived cells—that persist even in the immunocompromised mice we employed. Supporting this notion, tumor regression triggered by osimertinib has been associated with local accumulation of tumor-associated macrophages ^68^. Importantly, Bis3 exhibited in vivo activity comparable to that of its parental antibody combination, despite reduced antigen affinity of its anti-EGFR arm. This effect may be attributed to the bivalent architecture of Bis3 and the enhanced binding strength of its anti-MERTK arm, which appears to compensate for the faster dissociation rate of the anti-EGFR arm.

In summary, we initially observed that the upregulation of both GAS6 and its receptor, MERTK, coincided with the initiation of patient resistance to osimertinib, suggesting that the GAS6–MERTK axis provides a route for drug escape. This hypothesis was reinforced by the development of mAbs targeting GAS6 and MERTK, each of which partly delayed the emergence of TKI resistance. To streamline this approach, we next engineered a MERTK/EGFR bsAb, thereby eliminating the need for a separate anti-EGFR monoclonal. These findings not only establish the GAS6–MERTK pathway as a driver and therapeutic target of drug resistance, but also point to an alternative treatment strategy for patients with EGFR-mutant lung cancer, including exon-20 carriers: rather than relying on TKI monotherapy, which typically leads to resistance within about one year, combine the TKI with a bsAb that simultaneously targets MERTK and EGFR. In a clinical setting, such a combination would be expected to substantially delay, or even prevent, resistance while maintaining a favorable safety profile.

## Acknowledgments

We thank all current and past members of our laboratories, including Nishanth B. Nataraj, Aakanksha Verma and Ashish Noronha, for their kind help and insightful comments. This work was performed in the Marvin Tanner Laboratory for Research on Cancer. YY is the incumbent of the Harold and Zelda Goldenberg Professorial Chair in Molecular Cell Biology. Our studies were supported by the Israel Science Foundation, the European Research Council (ERC), the Israel Cancer Research Fund (ICRF), the Dr. Miriam and Sheldon G. Adelson Medical Research Foundation and the Medical Research Council, UK (grant number MR/w016176/1).

## Author contributions

Conceptualization, S.G. and Y.Y.; Methodology, S.G., B.S., A.SN., N.G., M.L., R.C., A.G. M.VD., D.R., L.R., M.Z., R.O., I.S.M., E.F.W., L.E.K., S.P., D.R., Y.A. and R.D.; Formal Analysis, S.G., B.S., A.SN., N.G., M.L., L.E.K.; Investigation, S.G., B.S., A.SN., N.G., M.L., L.E.K.; Resources, S.P., S.J.H., M.L., Y.Y.; Writing – Original Draft, S.G. and Y.Y.; Writing – Review, all authors; Supervision, E.R., S.J.H., M.L. and Y.Y.; Project Administration, Y.Y. Funding acquisition, S.J.H. and Y.Y.

## Inclusion and Diversity

We support inclusive, diverse, and equitable conduct of research.

## Materials and methods

### Cell cultures

PC9 cells were purchased from Sigma ((RRID:CVCL_B260)) and both H1975 ((RRID:CVCL_1511)) and HEK293T ((RRID:CVCL_0063)) cells were from ATCC. Cells were grown with either RPMI-1640 or DME medium supplemented with 10% fetal bovine serum (FBS) and Penicillin-streptomycin at 37°C under 5% CO_2_. Osimertinib (Cat. No.: HY-15772) and MRX-2843 (Cat. No.: HY-101549) were purchased from MedChemExpress. Recombinant cetuximab was expressed and produced in an Expi293™ (Thermo Fisher Scientific Cat# A14527) Expression System. The anti-GAS6 and anti-MERTK antibodies were generated using the hybridoma technology.

### Animal studies and PDX models

All studies were approved by the Weizmann Institute’s Animal Care and Use Committee (IACUC), which serves as the Institutional Review Board (IRB).CD1 nude mice were used for cell line xenograft studies, whereas NSG mice were used in experiments that tested PDX models. PC9 and H1975 cells (2.5 x 10^6^ per animal) were subcutaneously injected into the right flanks of 5–6-week-old female CD1 nude mice. Once tumors became palpable, the mice were randomized into treatment groups and administered TKIs via daily oral gavage or mAbs, twice weekly intraperitoneally. The PDX models, TM00193 and J000100672, were obtained from The Jackson Laboratory. Tumor fragments were implanted subcutaneously into the lower backs of 8–10-week-old male or female NSG mice (RRID:IMSR_JAX:005557). The procedure was performed under anesthesia, and the surgical site was closed using clips. When tumors reached approximately 350 mm³, mice were randomized into experimental groups as specified. Tumor volume was monitored twice weekly and calculated using the formula: (T1 × T2 × (T1 + T2)/2) / 2, where T1 and T2 are the tumor’s longest and shortest dimensions, respectively. Animals’ weights were determined once per week. Animals showing signs of illness were humanely sacrificed. The experimental endpoint was euthanasia by CO₂ administration.

### Generation of MERTK- and GAS6-overexpressing Cells

To generate PC9 cells overexpressing GAS6 or MERTK-eGFP we made use of lentiviral particles. These cells were co-transfected with second-generation packaging plasmids pMD2.G (RRID:Addgene_12259) and psPAX2 (RRID:Addgene_12260) along with lentiviral expression vectors encoding either MERTK-eGFP or GAS6. The resulting viral particles were harvested and used to transduce PC9 cells, which were then selected under puromycin to establish stably overexpressing cell lines.

### CRISPR/Cas9-mediated knockout of MERTK and GAS6

The CRISPR–Cas9 technology was used to knockout the MERTK and GAS6 genes. Target sites were selected adjacent to the Protospacer Adjacent Motif (PAM) sequence using the ENSEMBL database (Ensembl (RRID:SCR_002344)). A double-stranded break was introduced at the selected locations to disrupt the respective gene transcripts.

### Cell viability and proliferation assays

Cells (1 × 10³ per well) were seeded in 96-well plates. After 24 hours, they were treated with the indicated drugs for 72 hours. Following treatment, MTT solution (0.5 mg/ml) was added and incubated for 4 hours to allow for formazan crystal formation. DMSO was then added to dissolve the crystals, and absorbance was measured at 570 nm using a microplate reader. Alternatively, cells were seeded in 96-well plates and treated daily from Day 1 to Day 9, as indicated. On the designated days, cell proliferation was measured using crystal violet staining. Briefly, cells were fixed with ice-cold methanol for 20 minutes, followed by staining with 1% crystal violet in methanol for 20 minutes at room temperature. After washing and drying, SDS (2%) was added to solubilize the dye, and absorbance was measured at 490 nm.

### Colony formation and drug-tolerant persister (DTP) assays

For colony formation assays, cells (1,000 per well) were seeded in 6-well plates. For DTP assays, cells were seeded at full confluency in 6-well plates. Treatments were applied the following day, with media refreshed every 3 days. After 7–10 days, cells were fixed with ice-cold methanol for 20 minutes and stained with 1% crystal violet in methanol for another 20 minutes at room temperature. Colonies were scanned using an a Photo Scanner (Epson Perfection 4870). Images from five non-overlapping fields were captured using a light microscope (Olympus Corporation). Quantification was performed using ImageJ (ImageJ (RRID:SCR_003070)).

### Spheroid assays

Cells (1×10^3^) were seeded in a BME pre-coated 96-well plate and embedded in 5% BME medium containing drugs, as indicated. The experiments were stopped after 7 (PC9 cells) or 14 days (PC9-AZDR cells). Photos were captured using an OpTech IB4 microscope, in brightfield. Percentages of covered area were assessed using ImageJ.

### RNA isolation and real-time PCR

Total RNA was extracted using the RNeasy Mini Kit (QIAGEN), following the manufacturer’s instructions. cDNA synthesis was performed using the qScript cDNA Synthesis Kit (Quantabio). Real-time PCR was conducted using SYBR Green Master Mix (Applied Biosystems) and gene-specific primers (primer sequences provided separately). Reactions were run on a Real-Time PCR system (Applied Biosystems). GAPDH served as an internal control for normalization using the ΔCt method.

### Immunofluorescence analyses

Cells (1×10⁴ per well) were seeded on glass coverslips in a 24-well plate. After three days, cells were fixed with paraformaldehyde (4%; PFA) at room temperature for 15 minutes, followed by two washes with saline. Cells were then blocked and permeabilized for 60 minutes at room temperature using a solution containing 1% BSA and 0.3% Triton X-100. After blocking, cells were incubated overnight at 4°C with an anti-Ki67 primary antibody (Cell Signaling Technology Cat#9449; RRID:AB_2797703). The following day, cells were washed with saline containing 0.05% Tween-20 and incubated with an Alexa Fluor 488-conjugated secondary antibody. DAPI and Rhodamine B-conjugated phalloidin were used for nuclear and actin cytoskeleton staining, respectively. Images were captured using a Zeiss spinning disk confocal microscope and processed with Zeiss ZEN 3.1 software. ImageJ was used for signal quantification.

### Immunostaining and quantification of intracellular receptors

PC9 cells (5×10^4^) were treated with either mAb745 (10 μg/ml), cetuximab (10 μg/ml), or the bispecific antibody Bis3 (20 μg/ml). Treated cells were fixed with PFA (4%) for 20 minutes at the following time-points: 0h, 2h, 4h, and 8h. Fixed cells were subsequently permeabilized with 0,01% Triton-X100 for 5 minutes to enable intracellular labeling. Cells were blocked with a solution of PBS containing BSA (1%). Subsequently, immunofluorescence staining was performed using antibodies directed against EGFR (ab289889, 1:500) and MERTK (CST#4319S, 1:100). Cells were incubated in primary antibody for 60 minutes, washed 3 times in saline, and secondary antibodies specific to the mouse section of the anti-EGFR antibody and the rabbit section of MERTK were used. Coverslips were then mounted using the ProLong Gold Antifade Mountant (P36934). Images were taken using a Zeiss inverted spinning disk confocal equipped with a 60X objective. Several Z-Stack (1μm Z-axis) were taken for analysis. To quantify the internalized receptors, the middle section of the cells was selected from the Z-Stack, and for each channel the average signal in the cytosol of the cells was calculated. The resulting signal intensity was used as a thresholding value to remove the background cytosolic noise. The remaining spots were counted using the function “analyze particles”. To quantify and compare the different treatments, the average value of time zero of each treatment was used to normalize each sample.

### Immunoprecipitation and immunoblotting analyses

Cells were washed with saline and extracted on ice using a buffer containing 50 mM HEPES (pH 7.5), 10% glycerol, 150 mM NaCl, 1% Triton X-100, 1 mM EDTA, 1 mM EGTA, 10 mM NaF, 25 mM β-glycerophosphate, 0.1 mM sodium orthovanadate and a protease inhibitor cocktail. Lysates were centrifuged at 14,000xg for 20 minutes at 4°C, and the supernatants were collected for immunoprecipitation or immunoblotting. For immunoprecipitation, protein G beads were incubated overnight at 4°C with the indicated primary antibodies on a mechanical rotor. Beads were washed with ice-cold HNTG buffer (20 mM HEPES pH 7.5, 150 mM NaCl, 0.1% Triton X-100, 10% glycerol). Bound proteins were eluted by boiling in Laemmli buffer (Bio-Rad) and resolved using gel electrophoresis. This was followed by transfer to nitrocellulose membranes. Membranes were blocked in 3% BSA, incubated overnight at 4°C with primary antibodies, and then with HRP-conjugated secondary antibodies for 1 hour at room temperature. Blots were visualized using Clarity Western ECL Substrate (Bio-Rad) and imaged on a ChemiDoc Imaging System (Bio-Rad) and Image Lab v6.0.1 (RRID:SCR_014210).

### Sandwich ELISA

A custom sandwich ELISA was performed to assess the simultaneous binding of Bis3 to MERTK and EGFR. ELISA plates were coated overnight with the MERTK antigen, blocked with 3% BSA, followed by incubation with Bis3 for 60 minutes at room temperature. Next, the plate was incubated with an AVI-tagged soluble form of EGFR (SinoBiological Cat: 10001-H27H-B). Detection was carried out using Streptavidin-HRP, and absorbance was measured at 415 nm using a microplate reader.

### NanoBRET assays

HEK293T cells were seeded at a density of 500,000 cells per well into 6 well plates (Corning) and left to grow at 37°C/5% CO2 overnight. The next day, cells were transiently transfected with 500ug/well of EGFR and MERTK constructs, N-terminally tagged with NLuc (energy donor) or HaloTag (energy acceptor) to give a total of 1000ug cDNA per well using FuGENE HD (Promega Corporation) at a 3:1 DNA: reagent ratio in OptiMEM (Gibco) following the manufacturer’s instructions. The next day, cells were seeded onto white flat bottom 96 well microplates (Grenier Bio-One 655098) at a density of 40,000 cells per well. The next day, the medium was replaced with prewarmed assay buffer (HEPES-buffered saline solution supplemented with 0.1% protease-free bovine serum albumin (Sigma); termed HBS/0.1% BSA). Cells were then incubated with 0.2 μM membrane impermeable HaloTag AlexaFluor488 (Promega Corporation) substrate for 30 min at 37°C. Cells were then washed three times with HBS/0.15 BSA before being treated with the indicated growth factors. After incubation for 60 minutes at 37°C, the NanoGlo substrate furimazine (1:400 dilution) was added, and cells were further incubated for 5 minutes. Simultaneous fluorescence and luminescence emissions were then detected using a PHERAstar FSX plate reader (BMG LabTech), fitted with a bandpass filter to detect donor emission (NLuc emission) at 475 ± 30 nm and a bandpass filter to detect acceptor emission (HaloTag AlexaFluor488) at 535 ± 30 nm. BRET ratios were calculated by dividing fluorescence by luminescence emissions and baseline corrected to NLuc donor-only conditions.

### Proximity Ligation Assays (PLA)

Cells were plated on glass coverslips, fixed, permeabilized, and blocked using standard protocols. Primary antibodies were applied as indicated. The DuoLink In Situ PLA Detection Kit (Sigma-Aldrich) was used according to the manufacturer’s instructions. This included incubation with PLA probes, ligation, signal amplification, and detection. Images were acquired using a Zeiss LSM800 confocal microscope, and PLA signals per cell were quantified using ImageJ software.

### Surface Plasmon Resonance (SPR) Measurements

SPR was performed using the Biacore™ T200 system to determine the binding affinities of the bispecific antibody Bis3 and its parental antibodies—anti-MERTK (mAb745) and Cetuximab (CTX). Sensor Chip CM5 (Cytiva; Cat# BR100530) was activated using a freshly prepared solution of N-hydroxysuccinimide and 1-ethyl-3-(3-dimethylaminopropyl) carbodiimide to introduce reactive amine groups. MERTK and EGFR proteins were immobilized onto designated flow cells via covalent attachment. Unreacted groups were blocked using ethanolamine hydrochloride. Various concentrations of the antibodies were injected, and sensograms were recorded. After each cycle, the chip’s surface was regenerated using 2 mM NaOH. Data were analyzed using a steady-state affinity model of the Biacore T200 Evaluation Software.

### Drug synergy analysis

Cells (3×10³ per well) were seeded in 96-well plates and treated for three days with TKIs, either as monotherapies or in combinations, at varying concentrations. Thereafter, cells were fixed with ice-cold methanol for 20 minutes and stained with 1% crystal violet in methanol for another 20 minutes at room temperature. After staining, 2% SDS was used to solubilize the dye, and absorbance was measured at 595nm using a microplate reader. Cell viability was calculated relative to untreated controls. Drug synergy scores (osimertinib and MRX-2843) were quantified online using the Loewe model via SynergyFinder (https://github.com/IanevskiAleksandr/SynergyFinder#readme) (RRID:SCR_026127).

### Generation of monoclonal antibodies

Balb/c mice were immunized with the extracellular domain of MERTK (SinoBiological, Cat: 10298-H03H; 25 µg per injection). Blood samples were collected after the third and fifth immunizations to assess serum antibody titers. After seven total immunizations, including a final booster, mice were sacrificed, and splenocytes were harvested. These were fused with NSO myeloma cells to generate hybridomas, which were screened using ELISA and validated using immunoprecipitation. Positive clones were subcloned and selected. Anti-GAS6 antibodies were generated using a similar hybridoma approach. Monoclonal antibodies were purified from hybridoma supernatants using protein G affinity chromatography.

### Design, expression, and purification of recombinant antibodies

The heavy and light chain sequences of cetuximab were retrieved from public databases, while those of mAb745 were obtained from the hybridoma clones. DNA sequences encoding the variable regions were cloned into mammalian expression vectors containing human IgG1 and kappa light chain constant regions. For the design of the bispecific antibody, the EGFR-binding arm was formatted as a CrossMab (CH1–CL swap) to ensure correct heavy–light chain pairing. Next, heterodimerization of heavy chains was achieved through the introduction of mutations in the CH3 domains: Y349C/T366S/L368A/Y407V (EGFR arm) and S354C/T366W (MERTK arm). Antibodies were produced in Expi293 cells (ThermoFisher) using transient transfection and purified from the culture supernatant using protein G chromatography.

### Amino acid sequences of the four chains of Bis3 (the variable domain sequences are shown in Bold type)

i. Anti-MERTK (mAb745), heavy chain (‘knob’ mutations shown in green) MGWSCIILFLVATATGVHS**DVQIQESGPGLVKSSQSLSLTCTVTGYSITNDYAWNWIRQFPGN KLEWMGYISYNGNTFYNPSLKSRISITRDTSKNQFFLQLSSVTTEDTATYYCGRLLPVSYAM DYWGQGTSVTVSS**ASTKGPSVFPLAPSSKSTSGGTAALGCLVKDYFPEPVTVSWNSGALTSGVH TFPAVLQSSGLYSLSSVVTVPSSSLGTQTYICNVNHKPSNTKVDKRVEPKSCDKTHTCPPCPAPELLGGPSVFLFPPKPKDTLMISRTPEVTCVVVDVSHEDPEVKFNWYVDGVEVHNAKTKPREEQYNS TYRVVSVLTVLHQDWLNGKEYKCKVSNKALPAPIEKTISKAKGQPREPQVYTLPP**C**REEMTKNQ VSL**W**CLVKGFYPSDIAVEWESNGQPENNYKTTPPVLDSDGSFFLYSKLTVDKSRWQQGNVFSCS VMHEALHNHYTQKSLSLSPGK
ii. Anti-MERTK (mAb745) light chain MGWSCIILFLVATATGVHS**DVVMTQTPLSLPVSLGDQASISCRSSQSLVHSNGNTYLHWYLQ KPGQSPKLLIYKVSNRFSGVPDRFSGSGSGTDFTLKISRVEAEDLGVYFCYQSTHVPWTFGG GTKLEIK**RTVAAPSVFIFPPSDEQLKSGTASVVCLLNNFYPREAKVQWKVDNALQSGNSQESVT EQDSKDSTYSLSSTLTLSKADYEKHKVYACEVTHQGLSSPVTKSFNRGEC
iii. Anti-EGFR (cetuximab), heavy chain (‘hole’ mutations shown in green) MGWSCIILFLVATATGVHS**QVQLKQSGPGLVQPSQSLSITCTVSGFSLTNYGVHWVRQSPGK GLEWLGVIWSGGNTDYNTPFTSRLSINKDNSKSQVFFKMNSLQSNDTAIYYCARALTYYDY EFAYWGQGTLVTVSS**ASVAAPSVFIFPPSDEQLKSGTASVVCLLNNFYPREAKVQWKVDNALQ SGNSQESVTEQDSKDSTYSLSSTLTLSKADYEKHKVYACEVTHQGLSSPVTKSFNRGECDRSEPKSCDKTHTCPPCPAPELLGGPSVFLFPPKPKDTLMISRTPEVTCVVVDVSHEDPEVKFNWYVDGVEVHNAKTKPREEQYNSTYRVVSVLTVLHQDWLNGKEYKCKVSNKALPAPIEKTISKAKGQPREPQ V**C**TLPPSREEMTKNQVSL**S**C**A**VKGFYPSDIAVEWESNGQPENNYKTTPPVLDSDGSFFL**V**SKLTV DKSRWQQGNVFSCSVMHEALHNHYTQKSLSLSPGKSAWSHPQFEK
iv. Anti-EGFR (cetiximab), light chain, CrossMab MGWSCIILFLVATATGVHS**DILLTQSPVILSVSPGERVSFSCRASQSIGTNIHWYQQRTNGSPR LLIKYASESISGIPSRFSGSGSGTDFTLSINSVESEDIADYYCQQNNNWPTTFGAGTKLELK**SSASTKGPSVFPLAPSSKSTSGGTAALGCLVKDYFPEPVTVSWNSGALTSGVHTFPAVLQSSGLYSL SSVVTVPSSSLGTQTYICNVNHKPSNTKVDKKVEPKSC

### Bioinformatics analyses

The Kaplan–Meier Plotter tool (https://kmplot.com/analysis/) was used to evaluate the prognostic significance of MERTK expression in cancer patients. To perform the analysis, patient cohorts were selected based on available gene expression and survival data. The hazard ratio (HR) and corresponding p-values were automatically calculated by the tool and displayed on the survival plots. The data sources integrated into the Kaplan–Meier Plotter included several publicly available datasets from GEO and TCGA, covering lung adenocarcinoma patients from the following datasets (patient number) GSE102287 (n=66), GSE14814 (n=90), GSE157011 (n=235), GSE19188 (n=156), GSE29013 (n=55), GSE30219 (n=307), GSE31210 (n=246), GSE3141 (n=111), GSE31908 (n=40), GSE37745 (n=196), GSE43580 (n=150), GSE4573 (n=130), GSE50081 (n=181), GSE68465 (n=462), GSE77803 (n=156), GSE8894 (n=138) and TCGA (n=133). Single-cell RNA sequencing data were taken from a clinical trial, NCT03433469. Patients were treated with osimertinib, and the cohort was segregated into Treatment Naïve (TN), Residual Disease (RD), and Progressive Disease (PD). EGFR+ NSCLC patient samples were analyzed for differentially expressed genes (DEGs). Expression was compared between cells in the different time points using Wilcoxon’s test within the Seurat package, using R v4.2.1 (RRID:SCR_001905)).

### Statistical analyses

GraphPad Prism (version 10.6.0; GraphPad Prism (RRID:SCR_002798)) was used to perform statistical analyses. Sample numbers and other information (mean ± SEM, number of replicates, and specific statistical tests) are indicated in the respective figure legends. We employed one-way ANOVA followed by Dunnett’s multiple comparison test. The ImageJ software packages were used to perform data analysis. Statistical significance was denoted as follows: p<0.05 (*), p<0.01 (**), p<0.001 (***), and p<0.0001 (****). Differences were considered statistically significant when p<0.05. Unless otherwise indicated, all in vitro experiments were repeated twice.

