## Supplemental Information for "Co-targeting MERTK and EGFR with a Bispecific Antibody Overcomes Drug Resistance Across Mutations in Exons 19, 20, and 21"

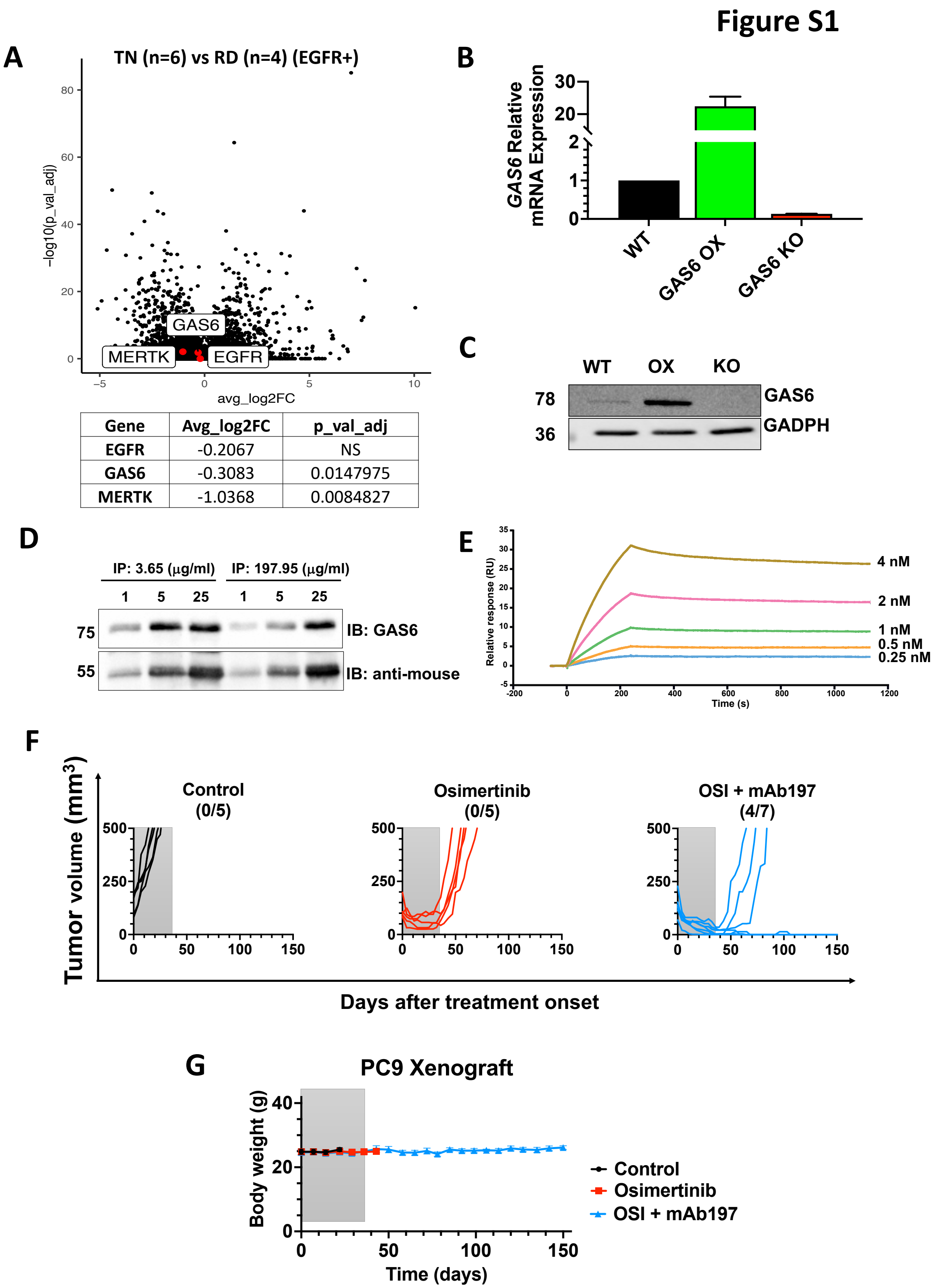


| Gene | Avg_log2FC | p_val_adj |
|---|---|---|
| EGFR | -0.2067 | NS |
| GAS6 | -0.3083 | 0.0147975 |
| MERTK | -1.0368 | 0.0084827 |

**Figure S1: Osimertinib treatment induces upregulation of GAS6 in patients and targeting GAS6 in an animal model delays the onset of resistance to osimertinib.** (**A**) Single-cell RNA sequencing data taken from NCT03433469 (EGFR+ NSCLC patients) were analyzed for differentially expressed genes (DEGs). Patients were treated with osimertinib, and segregated into Treatment Naïve (TN), Residual Disease (RD), and Progressive Disease (PD). The volcano plot presents DEGs that differ between TN (6 patients) and RD (4 patients). MERTK, GAS6, and EGFR are highlighted in red, and the average fold changes (log2FC) and P-values are presented in a table. (**B** and **C**) mRNA and protein extracts were isolated from the indicated genetically modified PC9 cells, and the expression of GAS6 was assayed using RT-PCR (B) and immunoblotting (C). GAPDH served as a gel loading control. (**D**) Anti-GAS6 mAbs derived from two hybridoma clones (clones 3 and 197) were tested at different concentrations (1, 5, and 25 μg/ml) in an immunoprecipitation assay. HEK293 cells were transfected with a GAS6 expression plasmid and culture supernatants were collected 48 hours post-transfection. The supernatant was subjected to immunoprecipitation using the specified antibodies and the resulting samples were analyzed by immunoblotting (IB) that used the indicated antibodies. (**E**) The binding affinity of the anti-GAS6 mAb197 was evaluated using Surface Plasmon Resonance (SPR) assays and the Biacore™ T200 system. Human recombinant GAS6 was utilized in this binding assay. The Sensor Chip was activated and bound to the recombinant GAS6 protein. Next, different concentrations of the anti-GAS6 mAb197 were passed through the specific channels. Sensograms were generated and the data were analyzed using a steady-state affinity model. (**F** and **G**) Shown are tumor volumes corresponding to individual mice comprising the three experimental arms presented in Figure 1D. Note that all treatments ended after 5 weeks (grey area). The numbers of surviving animals are shown in parentheses as a fraction of all animals tested in each experimental arm. Body weights were determined and the averages are shown in G. All experiments were repeated at least twice.

### Figure S2

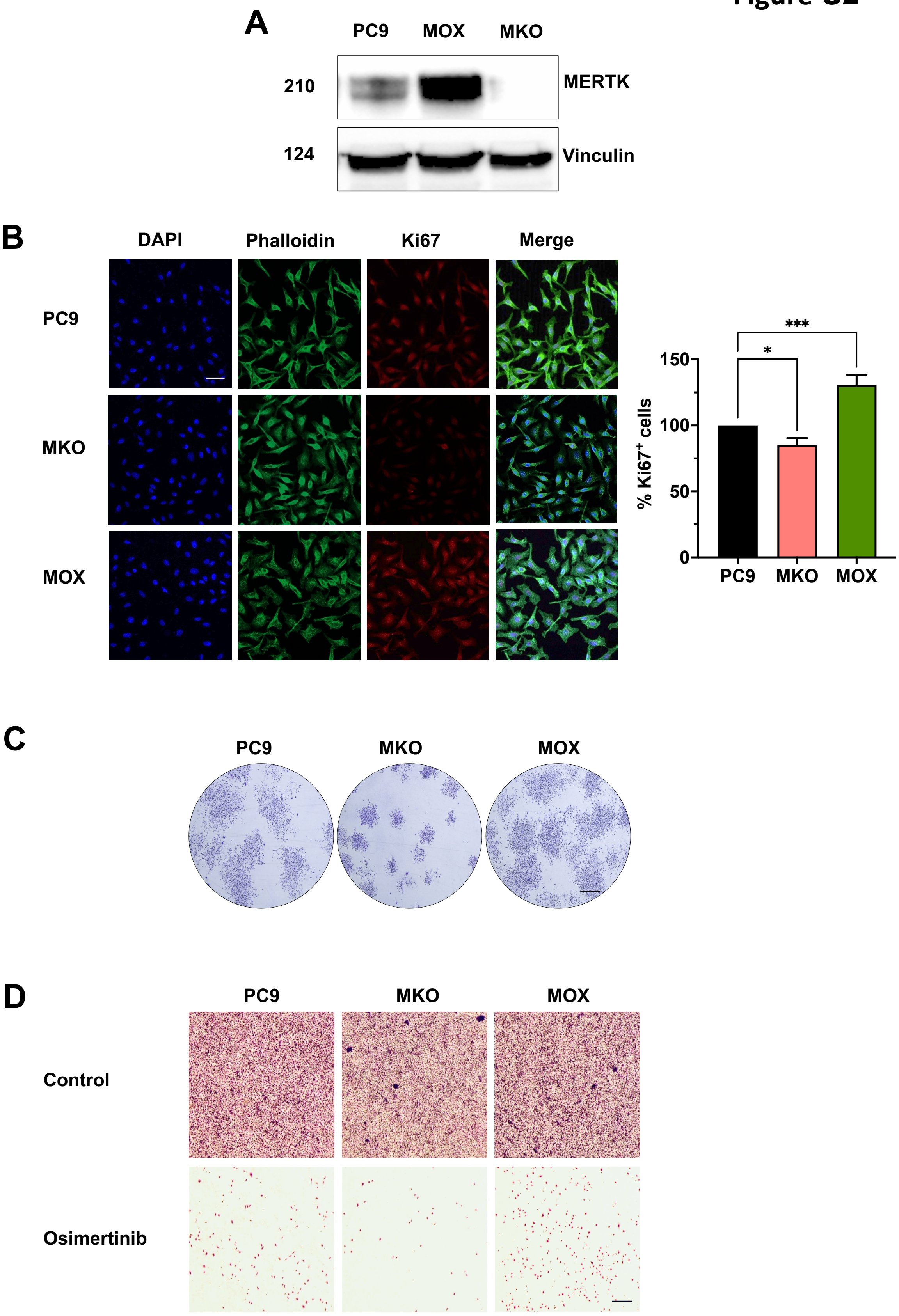
A
PC9
MOX
MKO
210
MERTK
124
Vinculin
B
DAPI
Phalloidin
Ki67
Merge
PC9
MKO
MOX
***
*
150
100
50
0
% Ki67+ cells
PC9
MKO
MOX
C
PC9
MKO
MOX
D
PC9
MKO
MOX
Control
Osimertinib

**Figure S2: MERTK is important for cell proliferation**. (**A**) Wild-type, MERTK knockout and MERTK overexpressing PC9 cells were cultured. Next, cell lysates were prepared for western blot analysis, as indicated, to test MERTK's expression level. Vinculin was considered a loading control. (**B**) Cells ($2x10^4$/per well) were seeded on coverslips with 1% serum and cultured for 5 days. Next, they were fixed and stained overnight with an anti-Ki67 antibody. Images were taken using a spinning disk confocal microscope and analyzed by ImageJ. The fractions of Ki67-positive cells, out of all DAPI-positive cells, were determined. The experiment was repeated twice. Scale bar, 4 µm. *, $p<0.05$; ***, $p<0.001$. (**C**) The indicated derivatives of PC9 cells ($1x10^3$ per well) were seeded on 6-well plates and incubated for 9 days. Thereafter, they were fixed and stained with crystal violet. Next, images were taken under a microscope and analyzed using ImageJ. Scale bar, 2 mm. See Figure 2D. (**D**) PC9 wild-type and the derivative cells were seeded in 6-well plates at high confluency. Next, they were treated, or not, for 9 days with osimertinib (0.3 µM). The media and drugs were refreshed once every 3 days. Finally, cells were fixed and stained with crystal violet. Images were taken from 5 non-overlapping fields with a light microscope and analyzed using ImageJ. Scale bar, 500 µm. All experiments were repeated at least twice. See Figure 2F.

### Figure S3

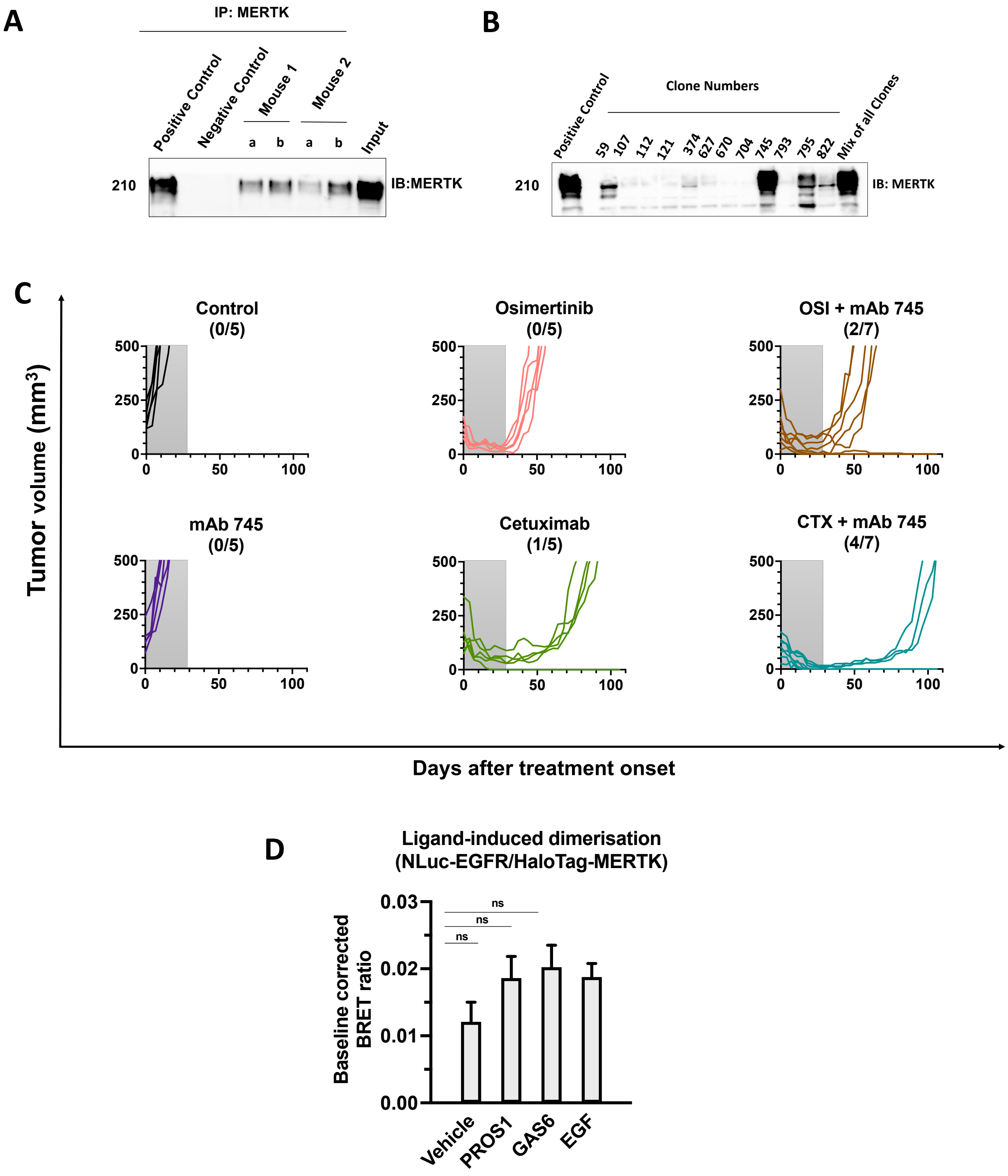


**Figure S3: Selection of anti-MERTK mAbs and in vivo xenograft trials**. (**A**) Human MERTK's extracellular domain was injected into mice at three-week intervals, for a total of six doses (25 µg/injection). Blood samples were collected in weeks 7 and 12, and the corresponding serum was used in an immunoprecipitation assay to evaluate the blood titre of anti-MERTK antibodies. PC9 cell lysate was used as the antigen source in this assay. As positive control, we used a commercially available anti-MERTK antibody. Pre-immunization serum was used as negative control. The immunoblot shows the levels of anti-MERTK antibodies present in the serum in week 7 (a) and in week 12 (b). (**B**) The second immunized animal was sacrificed in week 16 and the spleen was harvested for hybridoma generation. Approximately 1,000 hybridoma clones were established and initially screened for antibody titer using ELISA and immunoprecipitation assays. Selected positive clones were further evaluated by immunoprecipitation using PC9 cell lysate. A commercially available anti-MERTK antibody served as a positive control in this assay. (**C**) CD1-nu/nu mice bearing PC9 xenografts were randomized in groups of 5–7 animals and treated for 4 weeks with antibodies (200 µg, intraperitoneally, twice weekly) and/or daily with osimertinib (5 mg/kg, oral gavage). Tumor volumes were measured twice weekly, and body weight was measured once weekly. The grey-shaded area marks the treatment period. Changes in tumor volume ($mm^3$) from treatment onset are shown, with the number of mice (survived/total) indicated in brackets. (**D**) HEK293 cells were transfected with NLuc-tagged EGFR and Halo-tagged MERTK. Two days later, cells were incubated for 30 minutes with the HaloTag AlexaFluor488 substrate. Following three washes, cells were treated with the following ligands (each at 100 ng/ml): PROS1, GAS6 and EGF, for 60 minutes. The NanoGlo substrate furimazine was added and cells were further incubated for 5 minutes and luminescence and fluorescence emissions measured using a PHERAstar FSX and a 475 ± 15 nm bandpass filter (donor; NLuc emission) and 535 ± 15 nm bandpass filter (acceptor; HaloTag AlexaFluor488 emission). BRET ratios were calculated as the fluorescence emission/luminescence emission and normalized to the 'no acceptor' condition. Mixed-effects analysis and Dunnett's test for multiple comparisons were used. n.s, non significant. All experiments were repeated at least twice.

### Figure S4

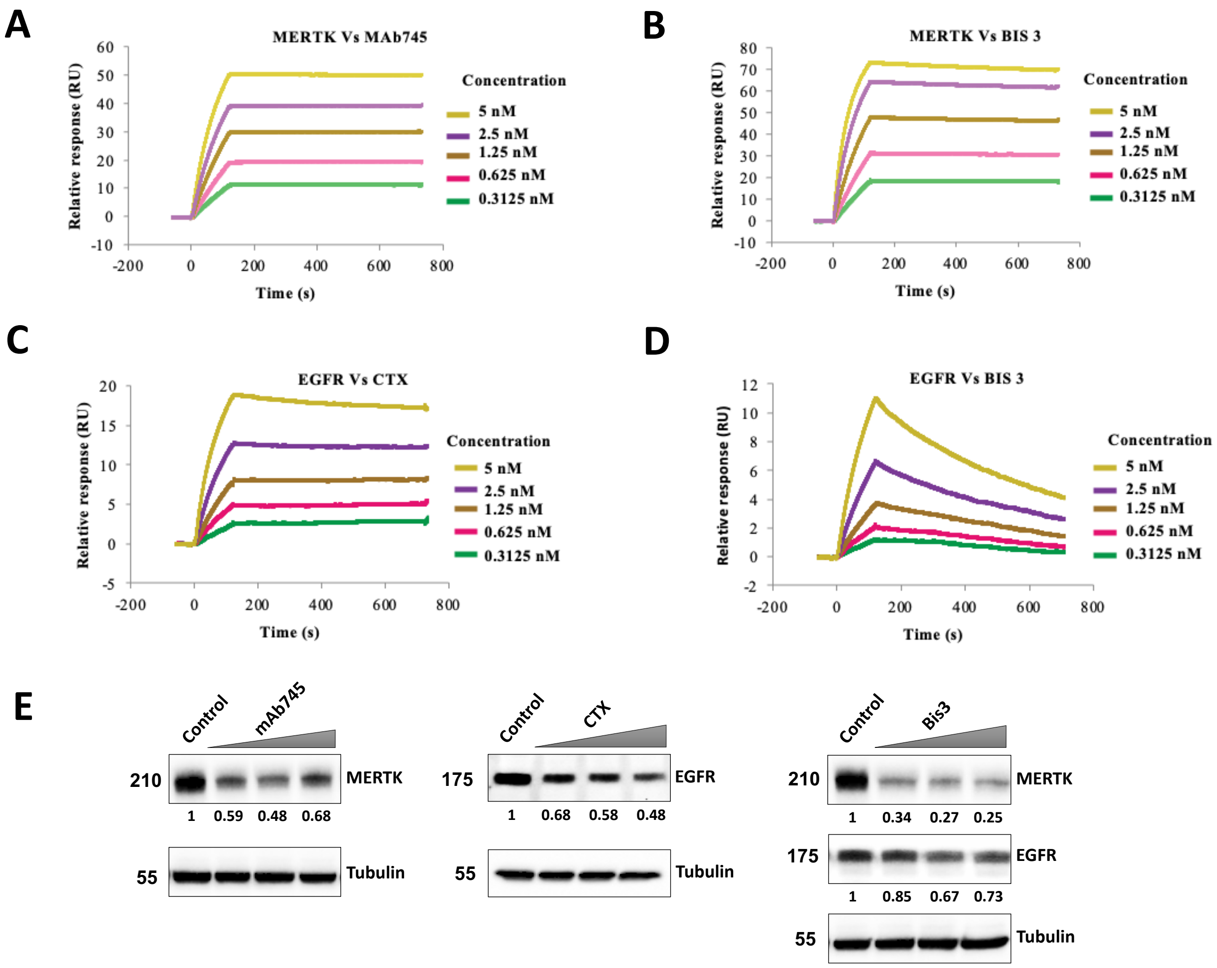


**Figure S4: Characterization of anti-MERTK mAbs and demonstration that Bis3 can induce degradation of both MERTK and EGFR.** (**A-D**) The binding affinities of the indicated antibodies towards EGFR and MERTK were assessed by analyzing the association and dissociation kinetics of antigen-antibody complexes using surface plasmon resonance (SPR; BIAcore 200 system). A recombinant MERTK protein was covalently immobilized on a sensor chip and various concentrations of mAb745 were flown over the surface for 2 minutes, prior to a 10-minute dissociation phase. Bis3 was tested on surfaces coated with both MERTK and EGFR in separate lanes, and the obtained signals were corrected by subtracting reference signals from empty lanes. The dissociation constant (Kd) was calculated as the ratio of the dissociation rate constant (koff) to the association rate constant (kon), and the values are summarized in Figure 4B. (**E**) PC9 cells were seeded in 6-well plates and subjected to overnight serum starvation. The following day, cells were treated for 48 hours with the indicated mAbs (5, 10 and 20 µg/ml), or Bis3 (10, 20 and 40 µg/ml), in serum-free medium. After treatment, cell lysates were collected and analyzed using an immunoblot assay that assessed receptor expression levels. Tubulin was used as a gel loading control. Estimated molecular weights are marked, along with densitometry-based signals of MERTK and EGFR (normalized to tubulin).

#### Figure S5

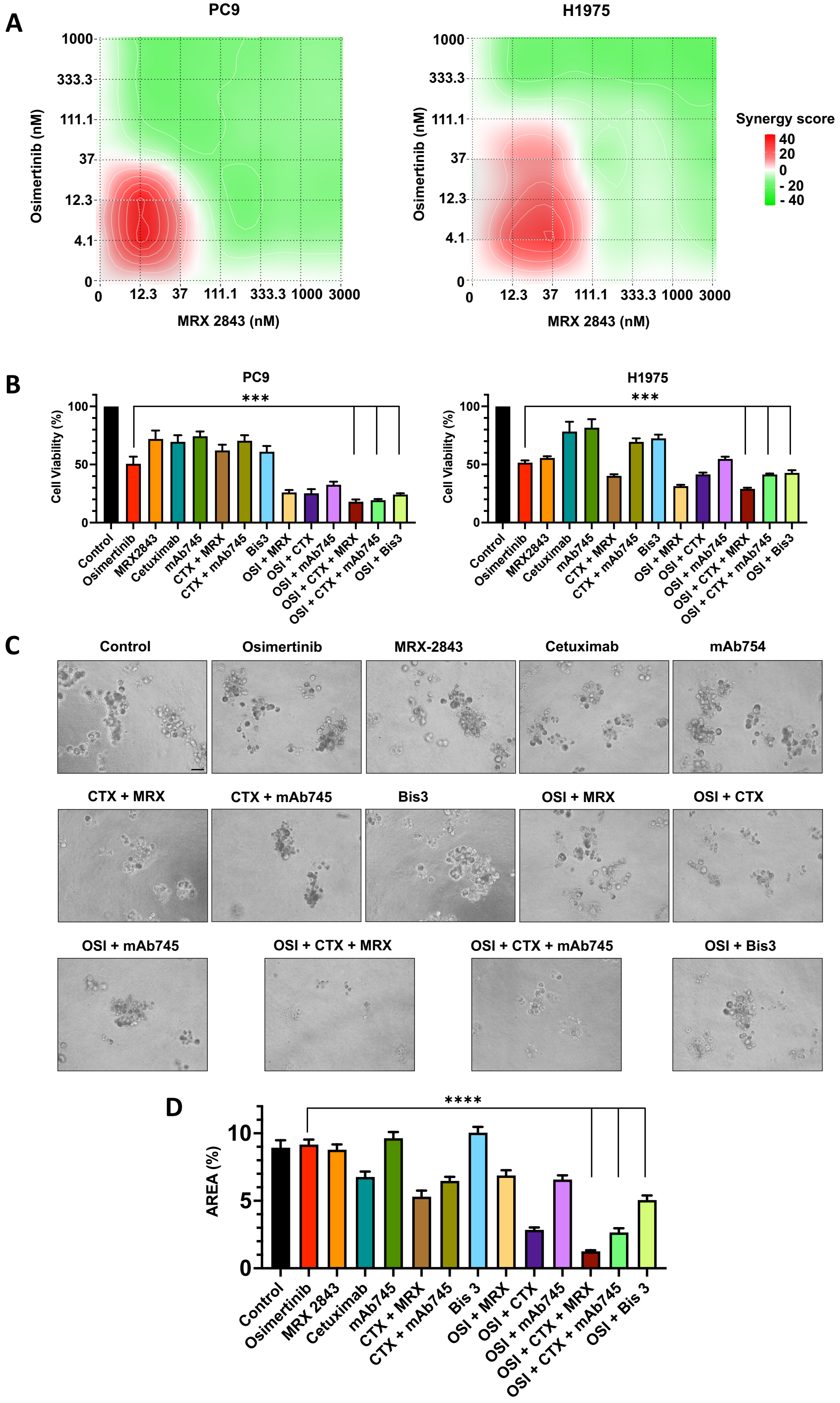
A
PC9
H1975
Osimertinib (nM)
MRX 2843 (nM)
1000
333.3
111.1
37
12.3
4.1
0
3000
Synergy score
40
20
0
- 20
- 40
B
PC9
H1975
Cell Viability (%)
***
Control
Osimertinib
MRX2843
Cetuximab
mAb745
CTX + MRX
CTX + mAb745
Bis3
OSI + MRX
OSI + CTX
OSI + mAb745
OSI + CTX + MRX
OSI + CTX + mAb745
OSI + Bis3
C
Control
Osimertinib
MRX-2843
Cetuximab
mAb754
CTX + MRX
CTX + mAb745
Bis3
OSI + MRX
OSI + CTX
OSI + mAb745
OSI + CTX + MRX
OSI + CTX + mAb745
OSI + Bis3
D
AREA (%)
****
Control
Osimertinib
MRX 2843
Cetuximab
mAb745
CTX + MRX
CTX + mAb745
Bis 3
OSI + MRX
OSI + CTX
OSI + mAb745
OSI + CTX + MRX
OSI + CTX + mAb745
OSI + Bis 3

**Figure S5: Pairs of either kinase inhibitors or mAbs, each co-targeting MERTK and EGFR inhibit a 3D model of osimertinib-resistant cells.** (**A**) PC9 and H1975 cells ($3\times10^3$) were seeded in 96-well plates. The next day, cells were treated with different concentrations of osimertinib and MRX-2843 in pairwise combinations for three days. Following fixation, cells were stained with crystal violet to identify viable cells. The synergy scores of both drugs were calculated based on the Loewe reference model, using SynergyFinder (see Methods). The red color indicates high synergy scores, implying cooperative EGFR-MERTK interactions. (**B**) PC9 and H1975 cells were seeded in 96-well plates. The next day, cells were treated for 72 hours as in Figure 5A. Cell viability was determined using the MTT colorimetric assay. The experiments were repeated twice using triplicates. Significance was calculated using one-way ANOVA, Dunnett's and Tukey's multiple comparison tests (**, $p < 0.01$). (**C** and **D**) An osimertinib-resistant derivative of PC9 cells (PC9-AZDR; $1x10^3$ cells) was seeded in BME pre-coated 96-well plates. The cells were embedded in 5% BME medium containing vehicle (CTR), the indicated mAbs (10 μg/ml), osimertinib (10 nM), MRX-2843 (300 nM), Bis3 (20 μg/ml) and the indicated drug combinations. The experiment was stopped after 14 days. Images of the spheroids were captured using an OpTech IB4 microscope (bright field; 20x magnification). The percentages of covered areas were assessed using ImageJ. Significance was calculated using one-way ANOVA, Dunnett's and Tukey's multiple comparison tests (**, $p < 0.01$; *** $p < 0.001$; ****, $p<0.0001$). The histograms represent the average of two independent experiments performed in duplicates. All experiments were repeated at least twice. Scale bar, 50 μm.

### Figure S6

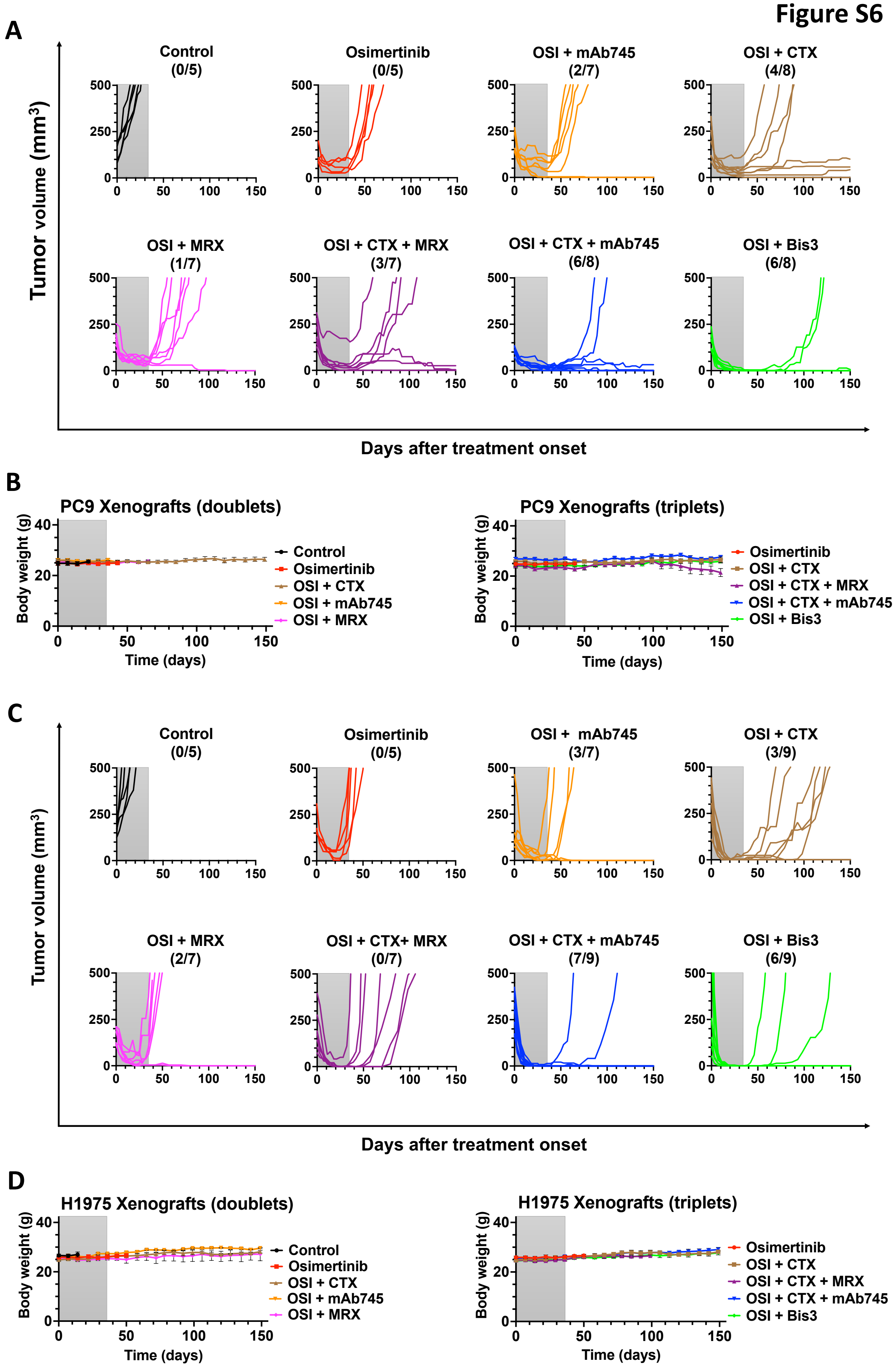
A
Tumor volume (mm3)
Control
(0/5)
Osimertinib
(0/5)
OSI + mAb745
(2/7)
OSI + CTX
(4/8)
OSI + MRX
(1/7)
OSI + CTX + MRX
(3/7)
OSI + CTX + mAb745
(6/8)
OSI + Bis3
(6/8)
0
250
500
0
50
100
150
Days after treatment onset
B
PC9 Xenografts (doublets)
Body weight (g)
0
20
40
Time (days)
Control
Osimertinib
OSI + CTX
OSI + mAb745
OSI + MRX
PC9 Xenografts (triplets)
Osimertinib
OSI + CTX
OSI + CTX + MRX
OSI + CTX + mAb745
OSI + Bis3
C
Tumor volume (mm3)
Control
(0/5)
Osimertinib
(0/5)
OSI + mAb745
(3/7)
OSI + CTX
(3/9)
OSI + MRX
(2/7)
OSI + CTX+ MRX
(0/7)
OSI + CTX + mAb745
(7/9)
OSI + Bis3
(6/9)
Days after treatment onset
D
H1975 Xenografts (doublets)
Body weight (g)
Time (days)
Control
Osimertinib
OSI + CTX
OSI + mAb745
OSI + MRX
H1975 Xenografts (triplets)
Osimertinib
OSI + CTX
OSI + CTX + MRX
OSI + CTX + mAb745
OSI + Bis3

**Figure S6: Combined inhibition of MERTK and EGFR shows efficacy and acceptable safety in cell line xenograft models.** (**A** and **B**) PC9 cells ($2.5x10^6$ per animal) were injected subcutaneously in the flanks of CD1-nu/nu mice. Once tumors became palpable, 5-9 mice were randomized and treated for four weeks with the indicated drugs. Tumor volumes (A) and body weights (B) were monitored. The drugs tested were either mAbs (100 µg/injection) or Bis3 (200 µg/injection), twice a week. In addition, where indicated, we used osimertinib (10 mg/kg) or MRX-2843 (20 mg/kg), which were orally administered daily. Tumor volumes were monitored twice a week, and animal body weight was determined once per week. Note that each line in A represents one animal but the data shown in B represent group averages. The number of mice surviving at the end of the experiment is shown in parentheses for each group. (**C** and **D**) The H1975 xenograft model was used and the results are presented essentially as described for the PC9 model in panels A and B.

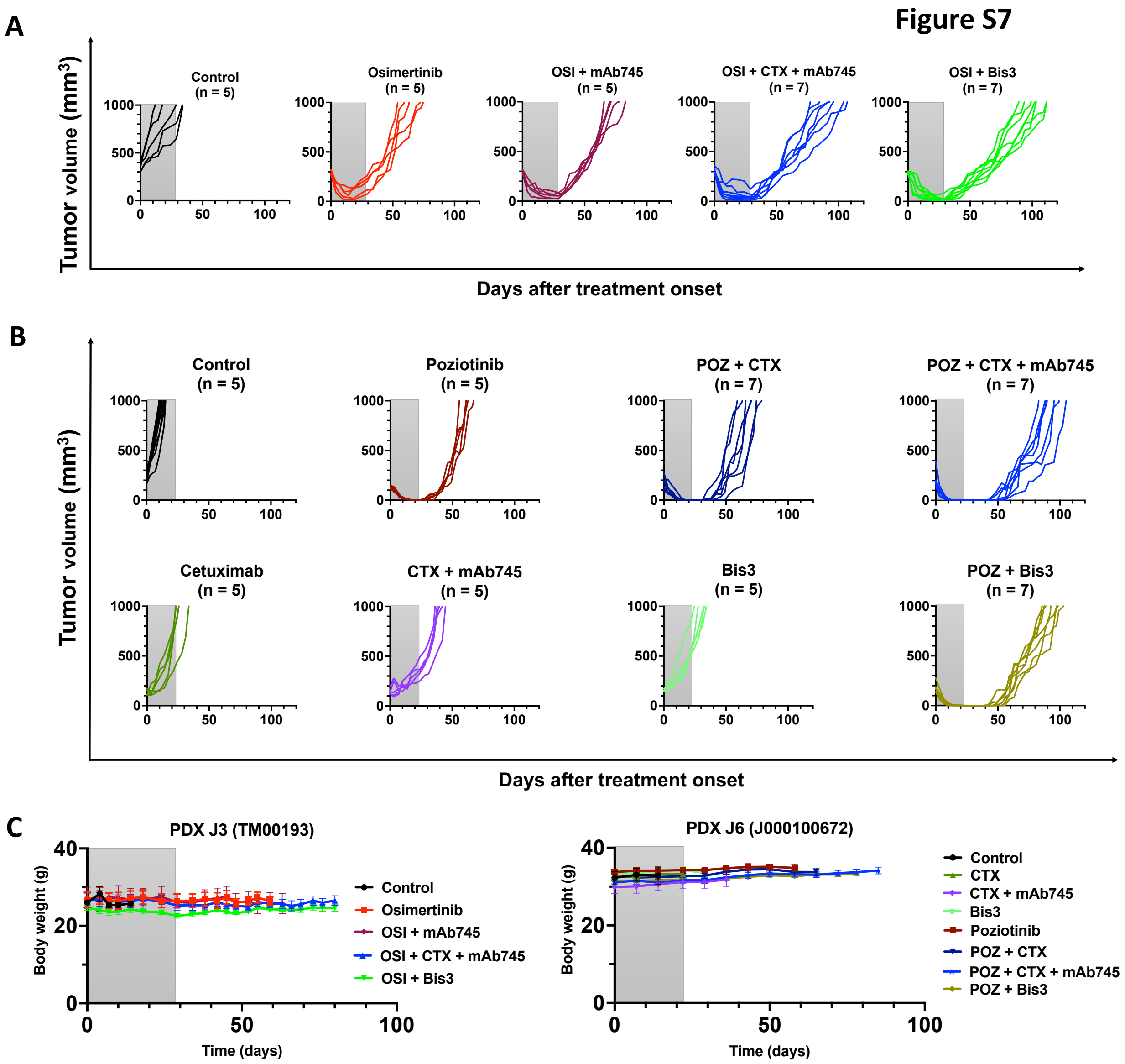


**Figure S7: The combination of Bis3 and either osimertinib or poziotinib displays a favorable safety profile and delays relapses in PDX models with exon-19 and exon-20 mutations.** (**A**) NSG mice bearing subcutaneous PDX TM00193 (EGFR Del19) tumors (~350 mm³) were randomized in groups of 5–9 animals per group. Mice were treated for 4 weeks with the indicated mAbs (100 µg/mouse; intraperitoneally, twice weekly), Bis 3 (200 µg/mouse, twice weekly), and/or osimertinib (10 mg/kg/day; orally, once per day). Tumor growth and body weight were monitored; 1000 mm³ was set as the sacrifice point. Changes in tumor volume from treatment onset are shown for each mouse. The number of animals in each group is indicated in brackets. (**B**) NSG mice bearing lung PDX6 (J000100672; exon-20 mutations) tumors (~300 mm³) were randomized (n = 5–7 per group) and treated for 3 weeks with antibodies and Bis3, as in A, and/or poziotinib (5 mg/kg; orally, once per day). Tumor growth and body weight were monitored with 1000 mm³ set as the sacrifice point. Changes in tumor volume from the onset of treatment are shown for each animal. The number of animals in each group is indicated in parentheses. (**C**) Body weight changes were monitored weekly throughout the experiments presented in A and B. Graphs show the average body weight for each treatment group, presented as mean ± SEM.
